# Enhanced reconstruction of Holocene bryophytes from sedimentary ancient DNA using bryophyte-specific primers

**DOI:** 10.1101/2025.11.15.688602

**Authors:** Bishnu Timislina, Dilli Prasad Rijal

## Abstract

The Arctic is warming rapidly, yet long-term data on how this affects bryophytes, key components of Arctic vegetation, remain limited. Sedimentary ancient DNA (sedaDNA) provides high-resolution taxonomic insights that can reveal vegetational responses to past environmental change, including functional responses via trait associations, but effective genomic tools for bryophyte sedaDNA are lacking. We applied bryophyte-specific primers targeting p6 loop in plastid DNA and compared their performance with bryophyte bycatch from vascular plant-specific primers. Using data from bryophyte-specific primers, we analyzed richness patterns across environmental gradients and assessed their potential for environmental reconstruction. The primers resolved nearly 60% of taxa to species level and over 80% to genus level, recovering 2.63 times more taxa with 1.48 times higher species-level resolution than vascular plant-specific primers. Taxonomic richness increased nonlinearly over time and along precipitation gradients, while exhibiting unimodal relationship with temperature and glacial activity. High-resolution data also enabled trait assignments for most bryophyte taxa, allowing temporal analyses of community development and functional dynamics. Overall, our results demonstrate that bryophyte-specific primers substantially improve bryophyte recovery from sedaDNA and, when combined with trait information, provide a powerful approach for reconstructing past bryophyte ecosystems and investigating functional ecological dynamics under long-term environmental change.

## Introduction

Bryophytes perform key ecological functions in regions with limited plant productivity, including the Arctic, where other vegetation is often scarce ^1^. They regulate and stabilize the soil temperature, retain moisture, and facilitate nutrient cycling. In addition to providing the local ecosystem services, they also play a significant role in moderating the impact of changing climate through high carbon sequestration ^2^. However, rapid warming is expected to severely affect the Arctic environment, reshaping vegetation structure, including that of bryophytes, and disrupting the ecological balances ^3^, making the future of bryophyte diversity in the Arctic concerning.

The rapid warming of the Arctic, known as Arctic amplification ^4^, is expected to increase the expansion of shrub cover, often called arctic greening ^5^, which may have cascading effects on bryophytes. As pioneer taxa, some bryophytes may take advantage of the new microclimate created beneath expanding shrubs, while others adapted to open, well-lit environments may be outcompeted and decline due to increased shading ^6^. Due to the lack of proper water retention mechanism, they are overall sensitive to change in temperature and precipitation which will potentially affect the diversity and composition ^7^. To understand the complexity of bryophyte responses to ongoing environmental change, long-term data are needed. Bryophytes’ responses to environmental change depend on their traits ^8^, and linking these traits to temporal data can help assess the impacts of change at the functional level. However, long term studies focusing on the impact of changing environment in bryophytes are lacking (but see, ref ^9^). Northern Fennoscandia experienced a significant change in glacial history, temperature and precipitation regime in the past ^10–13^. In this context, paleoecological records provide valuable insights about how bryophytes responded to changing environments in the past, providing a basis to predict their future responses.

Macrofossils are widely used as conventional tools to study temporal changes in bryophyte diversity and composition ^14–16^. Due to their high abundance, bryophytes are well represented in macrofossil records ^16,17^. Despite their availability and ability to discern taxonomy at higher resolution, macrofossils analysis may be limiting due to missing identifying features especially in the degraded macrofossils and the requirement of macrofossil identification skills ^16^, in the face of critically declining morpho-taxonomic expertise ^18^. In this context, sedimentary ancient DNA (*sed*aDNA) can be a potential alternative tool to study the past record of bryophytes. This tool has been proven to be effective in retrieving high resolution taxonomic data of vascular plants from the sediment depositions ^11,12,19,20^, often superior to traditional paleoecological methods ^19^. It can be leveraged to track temporal changes in composition and richness ^11^, reconstruct past ecosystems ^9,12^, track the species range shift ^21^ and reconstruct the past climate ^22^. As bryophytes have a narrow environmental niche and respond well to the environmental change ^23^, a similar approach can be applied to reconstruct the past environment. In addition, bryophytes are common bycatch in *seda*DNA analyses when targeting the vascular plants ^9,12^, hinting at the possibility of using *seda*DNA as a tool to study bryophyte palaeoecology.

Despite their potential, *seda*DNA of bryophytes have not been analyzed with high taxonomic resolution. Epp et al. ^24^ developed and tested metabarcoding primers specific to bryophytes suitable for degraded DNA. However, the results from the initial test were not promising ^25^. Steinthorsdottir ^26^ used modified reverse primer of the bryophyte-specific primer developed by Epp et al. ^24^ to analyze herbivore diets using fecal samples containing degraded environmental DNA, which resembles *seda*DNA, yielding promising results and hinting at the possibility of leveraging it for *seda*DNA studies. Here we test the use of bryophyte-specific primers as a metabarcoding tool to reconstruct past bryophyte communities. By applying it to lake sediments, we explore how bryophyte richness and community composition changed over time and how key traits varied through post-deglacial environmental shifts. This information will be crucial for understanding the change in bryophyte’s diversity and composition, especially under the current rapid environmental shifts occurring in the Arctic. Specifically, we aim to:

● Assess the effectiveness of bryophyte-specific metabarcoding primers in recovering bryophyte taxa,
● Analyze temporal trends in bryophyte richness and community composition, and
● Investigate temporal variation of traits in bryophytes.

## Methods

### Study area and sampling

The sediment cores used in this study were collected from Jøkelvatnet, a small lake (∼0.13 km²) located at 70.1725° N, 21.700833° E, and 158 m above sea level in the Sør-Tverrfjorddalen valley, northern Norway (Fig. 1). It is a distal glacial-fed lake that collects sediments reflective of upstream changes. The lake has a catchment area of about 11 km², drained by the Langfjordjøkelen ice cap from the south. The Tverrfjorddalen valley deglaciated completely ∼10 thousand calibrated year before present (ka) and started reformation approximately 4.1 ka ^27^. As the glacier is still active, it provides the opportunity of studying the change in vegetation accompanying the glacial change. Details of the study site and the sediment coring approach are provided in Wittmeier et al.^27^. We used 39 sediment samples taken from a 258 m long core (JØP-112; ^27^, collected in the GeoMicrobiology Lab at the Department of Earth Science, University of Bergen, by Rijal et al. ^11^. These samples were representative of the major shifts in the Holocene climate ^11^.

**Fig. 1:**
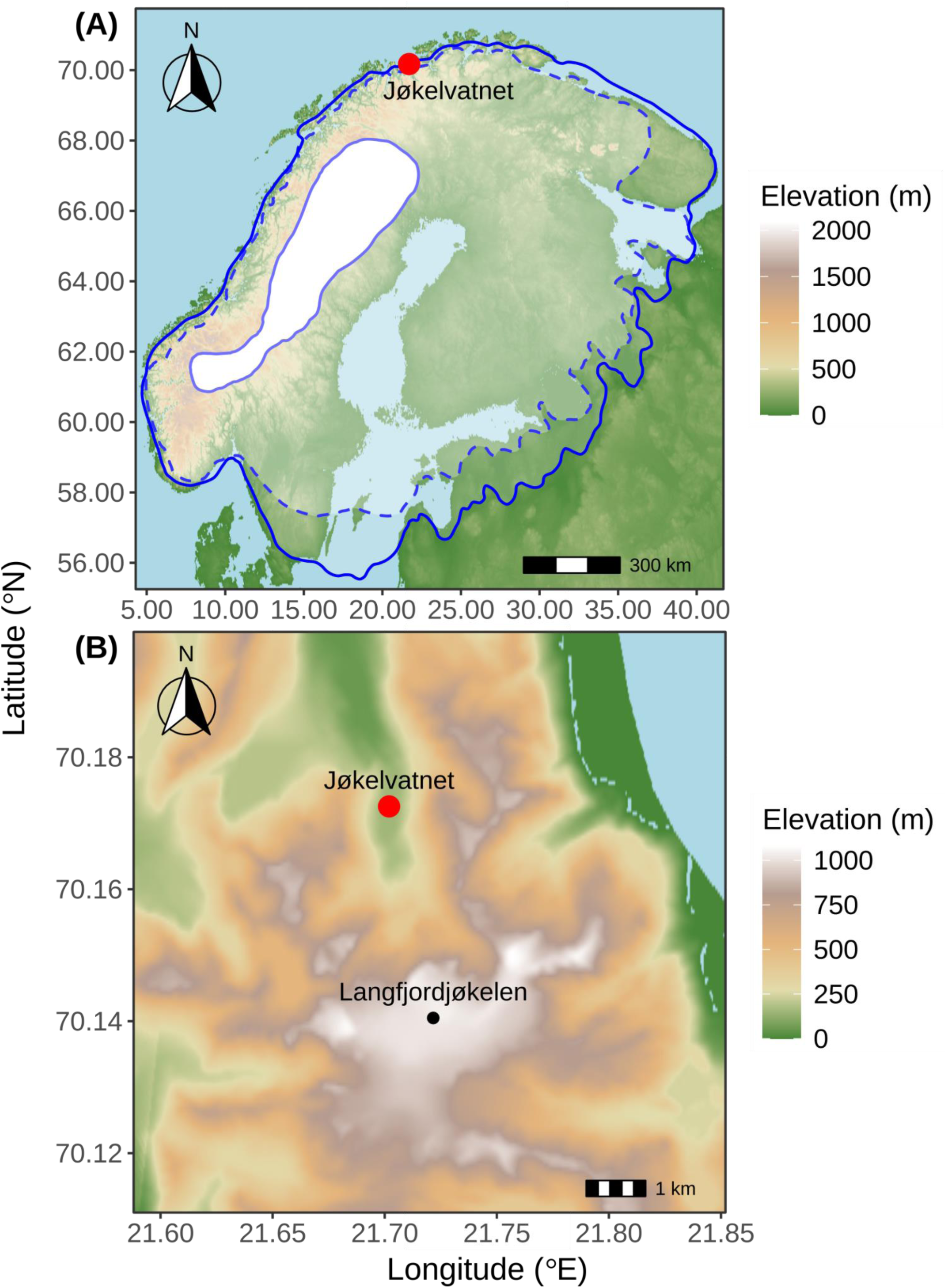
Study area map indicating (A) location of Jøkelvatnet along with extent of the Scandinavian ice sheet at 15, 14, and 10 ka, and (B) catchment area of Jøkelvatnet including Langfjordjøkelen glacier that drains into Jøkelvatnet. Note the scale differences of the axes.

### Molecular Laboratory Work

#### DNA extraction

We extracted DNA from 0.25-0.35 grams of the homogenized sediment samples using Qiagen’s DNeasy PowerSoil protocol with modifications involving addition of proteinase K and dithiothreitol followed by overnight incubation with constant rotation at 60 °C for effective lysis ^28^. Briefly, the samples were placed in a powerbid tube. The tube was centrifuged briefly, mixed with 750 μL of powerbid solution provided with the kit, and shaken in a fastprep machine for 20 seconds at the frequency of 4.0 m/s in two rounds, allowing the tube to cool. The mixture was then lysed with another reaction mixture containing 60 μL of solution C1, 25 μL of DTT, and 2 μL of proteinase K. The sample-lysis mixture was left for digestion of the organic materials present in the samples at the temperature of 56 °C overnight with constant rotation of 12 rpm. Following centrifugation, the suspension from the digested mixture was added with 250 μL of C2 solution, vortexed, and incubated at 2–8 °C for 10 min. It was then centrifuged to remove any precipitate, and the supernatant was added with 250 μL of C3 solution and incubated in cold as above. The supernatant was then treated with 1400 μL of C4 solution and ran through the spin column to retain the DNA. The column was then eluted with 65 μL of TET buffer twice to retain about 130 μL of DNA.

### *seda*DNA amplification and sequencing

For PCR multiplexing, we used uniquely tagged dual primer sequences consisting of 8 to 9 base pairs modified from Taberlet et al. ^29^ (Supplementary Table 1). The P6 loop of the *trn*L intron in plastid DNA was targeted for the amplification of bryophyte sequences (40-100 bp long) using a set of primers originally developed by Epp et al. ^24^ and later modified by Steinthorsdottir ^26^, in which a degenerate base (Y) was introduced at the fifth position of the reverse primer (CCATYGAGTCTCTGCACC), while the forward primer (GATTCAGGGAAACTTAGGTTG) remained unchanged. The PCR reactions were repeated eight times to maximize detection and minimize potential PCR biases. We used 8 controls including seven negatives consisting of extraction, DNA plating, PCR and post-PCR controls and one positive control with each of the replicates making a total of 376 PCR sets. The PCR mixture consisted of 4 μL of 10x Gold buffer, 4 μL of 2.5 mM MgCl2, 0.32 μL of 160 ng/μL of bovine serum albumin, 3.2 μL 0.2 mM of dNTPs, 0.32 μL of 1.6 U/μL of AmpliTaq Gold and 2 μL 0.1 mM of each of forward and reverse primer (total 4μL) and 4 μL template DNA and the final reaction volume was made to 40 μL with molecular grade water. Post-PCR negative and positive controls were introduced in the post-PCR lab. The thermocycle consisted of 10 minutes of initial denaturation followed by 45 cycles of denaturation for 30 seconds at 94 °C, annealing for 30 s at 55 °C, extension at 68 °C and the final extension for 10 minutes at 72°C. All the lab work was conducted in a dedicated ancient DNA laboratory of the Arctic University Museum of Norway.

We performed gel electrophoresis to check the presence of a precise amplicon in the PCR product. 10 μL of each of the PCR products were pooled into a tube and cleaned to remove the primer dimer, enzymes, or other impurities contained in the amplicon pool using MinElute PCR Purification Kit by Qiagen following manufacturer’s guidelines. In brief, we took 500 μL of pooled PCR product in a 5 ml tube and mixed it with 2500 μL of PB buffer and 50 μL of sodium acetate. Then 720 μL of the mixture was transferred to 4 spin columns placed in a 2 ml collection tube and centrifuged for 1 minute until the mixture was completely run through the spin column, followed by dry spin. Alcohol-based PE buffer was added to the volume of 720 μL and centrifuged, followed by another dry spin to remove the traces of alcohol in the spin column. It was then eluted with 20 μL of resuspension buffer twice to retain 40 μL of the cleaned pool of amplicons. The concentration of amplicons in the cleaned pool was estimated with Qubit before proceeding with library preparation. The cleaned pool of amplicons was subjected to end repair using the Illumina TruSeq DNA PCR-Free ’LS’ protocol, followed by purification with MinElute columns by QIAGEN, a modification adapted for *seda*DNA. This was followed by A-tailing and ligation of Illumina adapters under strict ancient DNA (aDNA) lab conditions, including the use of 80% ethanol and water from the aDNA facility, along with a negative control. Ligated fragments were purified using SPB (magnetic beads coated with a silica-based surface) beads with two rounds of 80% ethanol washes, and libraries were validated through Qubit quantification and qPCR. Sequencing was done in Nextseq (2x150) from Illumina at the Genomics Support Centre Tromsø, UiT-The Arctic University of Norway.

### Bioinformatics

The bioinformatics analysis was performed using OBITools4 ^30^. Forward and reverse fastq sequences overlapping with a minimum of 10 base pairs and having 80% similarly were paired using *obipairing* function, and the un-paired sequences were removed. The aligned sequences were filtered using the *obigrep* and were assigned to the corresponding samples using the *obimultiplex* function. The identical sequences collapsed to unique sequences and their numbers were counted using the *obiannotate*. The sequence tag and the rare sequence variants having less than 10 reads were removed. Similarly, the shorter fragments of DNA having a length less than 40 bp were also removed. The sequences that were not variants of another sequence with a count greater than 5% of their count were removed as chimeric sequences using *obiclean*. The reference library was created by *in silico* PCR of the EMBL sequences, allowing a maximum of three mismatches, using the *obipcr* function. Two sets of libraries were prepared, one with a global bryophyte taxa and the other with a subset of the bryophyte taxa found in Europe. The sequences were matched to the reference libraries, and taxonomic assignment was performed using the *obitag* function. Only sequences with a 100% match to either reference library were retained for downstream analysis (Supplementary Table 2). Taxonomic assignments matching the global bryophyte library were further verified by manual BLAST searches against the NCBI database. The number of PCR repeats of particular taxa were retained for further analysis (Supplementary Table 3). The distribution of the resulting taxa were manually checked using the database on Norwegian Biodiversity Information Centre (https://artsdatabanken.no/) and Global Biodiversity Information Facility (https://www.gbif.org/).

### Statistical analysis

Hierarchical cluster analysis was performed using a stratigraphically constrained incremental sum of squares (CONISS) method ^31^ on the proportion of PCR repeats using Bray-Curtis distance in the *Rioja* package ^32^. The significant number of clusters was determined using a broken stick model. We calculated taxonomic richness as the total taxa present within a sample. Bryophyte richness captured by vascular-plant primers for the samples used in this study was calculated from the published bycatch data ^11,12^. To compare the temporal trend in richness captured by bryophyte and vascular plant-specific primers, we fitted a generalized additive model (GAM) with richness as the response and sample age as the predictor variables. Using the mgcv package ^33^, we evaluated various distributions for modeling count data. The final model was based on *Tweedie* distribution as it provided better model fit (Supplementary Figs. 1A-B) compared to *Poisson* and negative binomial distributions. We considered titanium release values from X-ray fluorescence analysis ^27^ (Supplementary Fig. 6A, Table 6) as a proxy for glacial activity and modelled its impact on taxonomic richness (Supplementary Fig. 2). Paleoclimate data retrieved from CHELSA-TRACE21k ^34^ (Supplementary Fig. 6BC, Table 6) were used to evaluate the effects of mean temperature and precipitation of the warmest quarter on bryophyte richness, as well as their combined effect, using a GAM with a *Gaussian* distribution (Supplementary Fig. 3). Prior to modeling, we assessed collinearity between the two predictors, which was low. The estimated concurvity of the fitted model, which measures nonlinear dependency among the smooth terms ^33^ was also low for interaction between the two covariates. DHARMa package ^35^ was used to test the compliance of the model fitting.

To compare the taxonomic composition during different periods of glacial activity, we first created a distance matrix using the Bray-Curtis method, based on the number of PCR repeats per taxon. Non-metric multidimensional scaling (NMDS) analysis was then performed to examine the taxonomic composition during the early high (∼ 10.4 to ∼ 9.6 ka), middle low (9.6 to 4.2 ka), and late high (4.2 ka to present) glacial activities using the vegan package ^36^. Permutational multivariate analysis of variance (PERMANOVA) was conducted with 999 permutations using the *adonis2* function to statistically test differences in taxonomic composition of bryophytes during different glacial activity periods. All the analyses were performed in R ^37^.

### Trait reconstruction

We used the bryophyte traits data ^38^ to reconstruct the temporal change in bryophyte communities regarding life form and emergence of different taxa adapted to differential environmental conditions. Ecological traits measured in the ordinal scale of 1-9 were regrouped to four levels. For example, the traits related to light on the scale of 1 to 9 were regrouped to shade (1, 2 and 3), moderate shade (4 and 5), moderate light (6 and 7) and light (8 and 9) to ease plotting and interpretations. Biome-related traits such as arctic-montane, boreo-arctic montane, and wide-boreal, which include overlapping bryophyte species found in Arctic regions, were regrouped into boreo-arctic montane. Similarly, wide-temperate and temperate categories were regrouped into temperate (Supplementary Table 7). The first appearance date (FAD) of each taxon was recorded as the earliest date the corresponding taxon was detected among samples (Supplementary Table 2), and it was evaluated for its association with bryophyte traits. Changes in the proportion of bryophyte traits were visualized using area plots showing the proportion of PCR repeats associated with each trait over time.

## Results

### Bryophyte-specific primers are effective in retrieving high resolution taxonomy

After quality filtering, we recovered 12,176,846 reads, of which 10,331,251 reads matched 100% to bryophyte sequences in the EMBL reference library, either from the global dataset or from a subset limited to European taxa. The matched sequences yielded 309 amplicon sequence variants (ASVs, Supplementary Table 2). The bryophyte-specific primer produced significantly longer amplicons compared to vascular plant-specific primers (Fig. 2A, *p* < 0.0001, median = 53 vs. 18 bp, Fig. 2A, Supplementary Table 2). The taxonomic richness captured by bryophyte-specific primer was also significantly higher compared to the bycatch data of vascular plant-specific primer (*p* < 0.0001, mean ± SD = 50.94 ± 35.84 vs. 14.62 ± 9.97, Fig. 2B). Similarly, GC content in the amplicon produced by bryophyte-specific primer was higher than that for vascular plant-specific primers (mean ± SD = 24 ± 4.46 vs.15.39 ± 10.54, Supplementary Fig. 4, Supplementary Table 4). The ASVs were assigned to 163 taxa, of which majority (65%) belonged to class Bryopsida followed by Jungermaniopsida (28%), Polytrichopsida (3%) and other classes (Supplementary Figs. 4A-D). Overall, bryophyte-specific primers detected a 2.63 times higher number of taxa compared to bycatch detections. The taxonomic resolution achieved using the bryophyte-specific primer was 1.48 times higher at the species level, and 1.36 times at the genus level compared to that obtained using the vascular plant-specific primer (Fig. 2C). In contrast, the bycatch data from the vascular plant-specific primers included a greater number of taxa identified only at coarser taxonomic ranks, such as family or above (Fig. 2C).

**Fig. 2:**
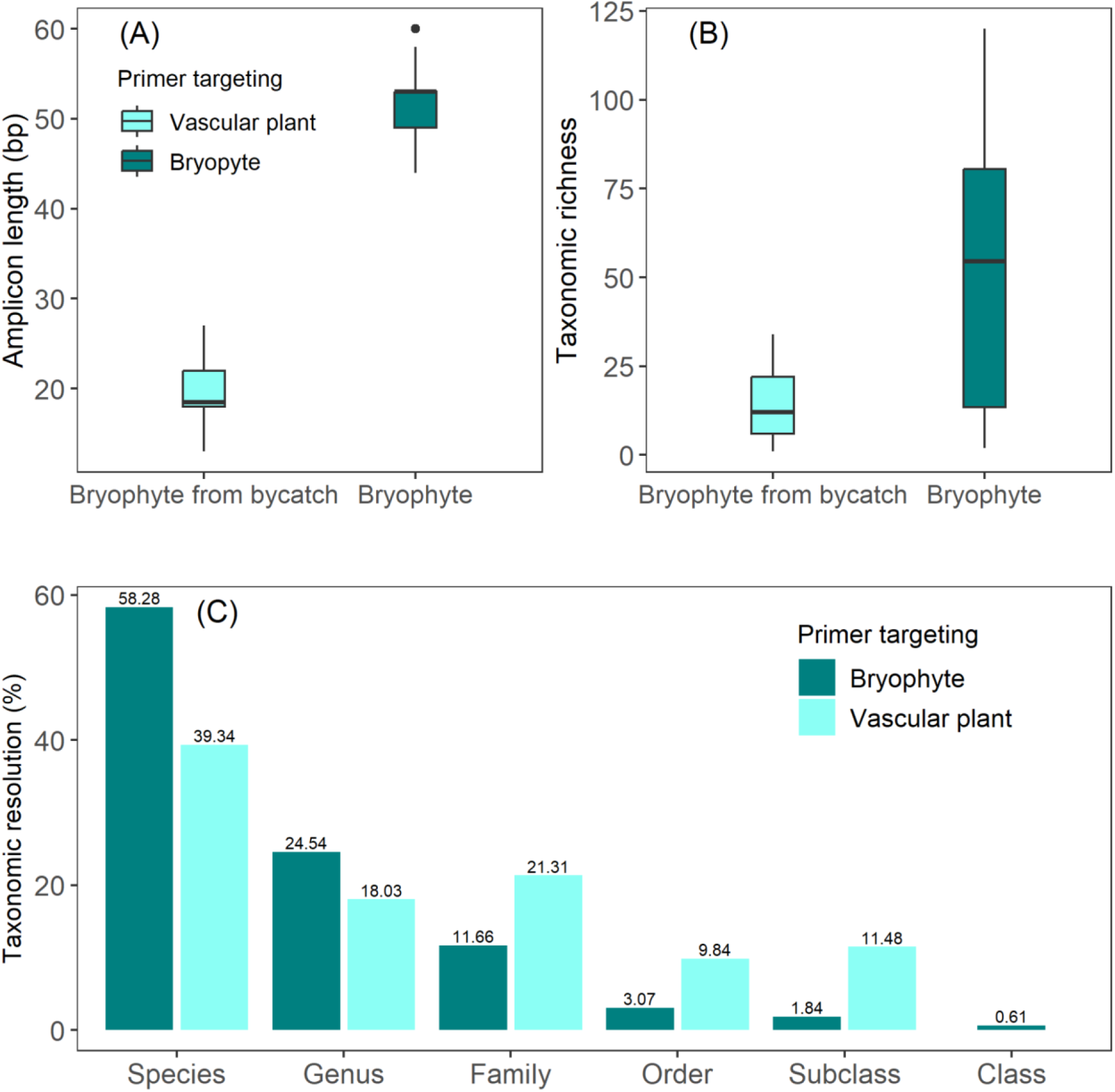
Comparison of the performance of bryophyte and vascular plant-specific primers. A) Difference in the amplicon lengths, B) Comparison of taxonomic richness, and C) Comparison of the taxonomic resolution.

### Temporal dynamics of bryophyte richness

Bryophyte richness increased over time following a complex nonlinear trend (edf = 4.567, *F* = 32.84, p < 0.001). The taxonomic richness steadily increased until ∼ 8 ka, followed by exponential increase until ∼ 4 ka. There was a subtle decrease in richness from ∼ 4 to 2 ka and then increased until present (Fig. 3A). However, the richness captured by vascular plant-specific primer increased continuously over time but remained always lower than that captured by bryophyte-specific primer (Fig. 3A).

**Fig. 3:**
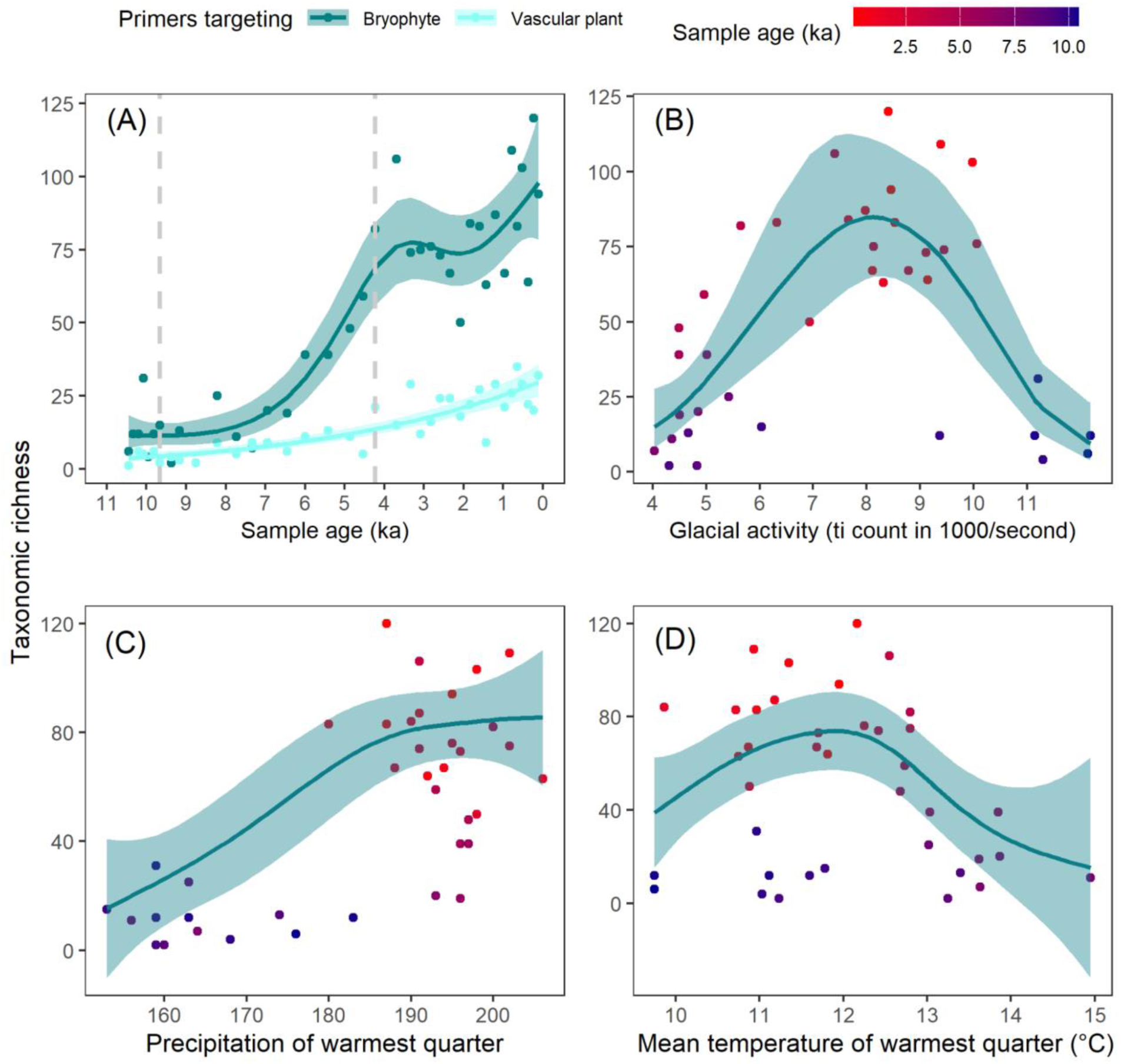
Taxonomic richness patterns of bryophytes A) over time captured by the bryophyte and vascular plant-specific primers (two vertical dashed lines enclose period of low glacial activity), B) with glacial activity measured as titanium release count in X-ray fluorescence analysis, C) with mean temperature of the warmest quarter, and D) with precipitation of the warmest quarter. The color gradient indicates the sample age with darker blue representing older and red younger samples.

### Richness trends and environmental correlates

Glacier activity had a unimodal relationship with the richness (*effective df (edf)* = 3.568, *F* = 10.54, *p* < 0.001, Fig. 3B). The model explained 57% variation in bryophyte richness (R²_adj_ = 0.56). Bryophyte richness increased with the mean precipitation of the warmest quarter, following a weakly nonlinear relationship (*edf* = 2.3, *F* = 17.258, *p* < 0.001). The pattern was characterized by low richness at low precipitation, a plateau or slight decline at intermediate values, and higher richness at high precipitation levels approaching an asymptote (Fig. 3C). However, richness exhibited an approximately unimodal relationship with the mean temperature of the warmest quarter (*edf* = 3.35, *F* = 5.2, *p* = 0.02), peaking at intermediate temperatures (∼12 °C) and declining toward both lower and higher temperature extremes (Fig. 3D). The mean temperature and precipitation of the warmest quarter had a significant interactive effect on bryophyte taxonomic richness (*edf* = 1, *F* = 5.91, *p* = 0.021). The GAM interaction model explained 73.6% of the variation in bryophyte richness (R²_adj_ = 0.68).

### Changes in bryophyte community composition

CONISS identified three distinct vegetation clusters based on taxonomic composition (Supplementary Fig. 6). The earliest vegetation zone covered the time from 10.45-9.37 ka, the second from 9.37 to 5.99 ka, and the third from 5.99 to present (see Supplementary text). This clustering roughly corresponded with the glacial activities in the catchment (Supplementary Fig. 7A).

NMDS analysis revealed three taxonomically distinct clusters corresponding to periods of early high (∼10.4 – 9.6 ka), middle low (9.6 – 4.2 ka), and late high (∼ 4.2 ka to present) glacial activity, with significantly different composition among the clusters (*F* = 9.00, *p* = 0.001; Fig. 4). The high glacial activities increased the variation in bryophyte community composition. Whereas there was little variation in the bryophyte composition during the low glacial activity period. The bryophyte community during the early high glacial period was composed of markedly different taxa, whereas communities during the middle low and late high glacial periods were more similar to each other (Fig. 4).

**Fig. 4:**
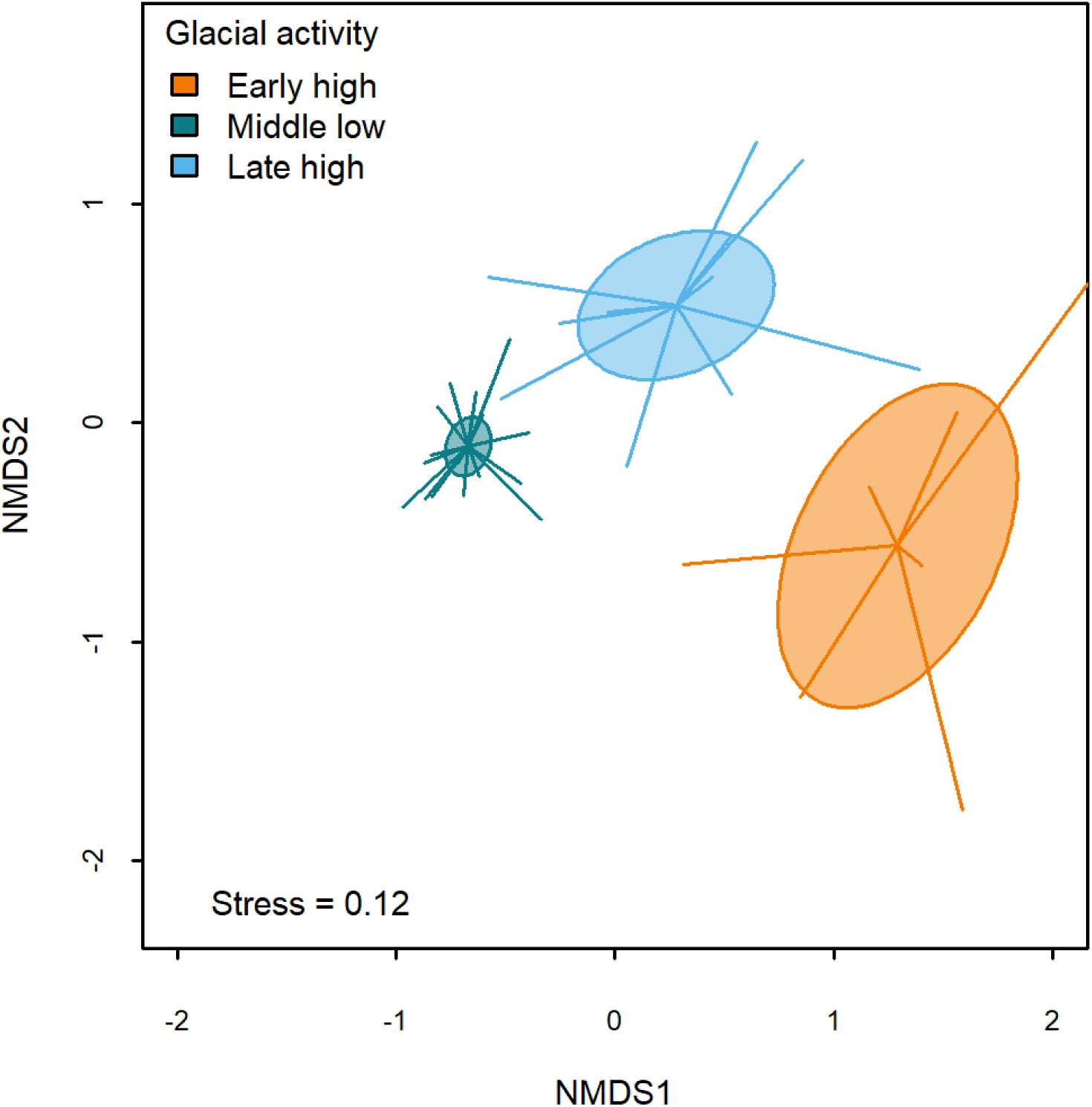
Bryophyte composition during early high, middle low and late high glacial activities based on the proportion of PCR replicates as the semi-quantitative abundance metrics. The size of the ellipses indicates compositional variation or dispersion.

### Trait variation through time

The r-selected taxa, which are composed of pioneer colonizers, dominated the post glacial phase from around 10.4-9.6 ka. As the glacier activity reduced, forming a more stable environment (Supplementary Fig. 7A), K-selected taxa were more abundant during the middle period with low glacial activity (9.5-4.2 ka), followed by an increase in r-selected taxa with the onset of increased glacial activity (Fig. 5A). There was also a gradual increase in turf life form though the proportion of cushion remained almost similar throughout. Weft life form peaked at ∼ 8 ka (Fig. 5B). Early Holocene taxa were characteristics of the boreo-arctic montane biome, the abundance of which continuously increased together with taxa of boreal montane biome until present; the taxa adapted to temperate environments peaked at around 7.5 ka (Fig. 5C). During the early Holocene, taxa adapted to high light conditions were abundant. The proportion of taxa adapted to moderate shade peaked between ∼8 and 6 ka, and the taxa adapted to light increased continuously after ∼ 8.5 ka (Fig. 5D). At ∼9 ka, taxa adapted to moderate nutrient levels were abundant, but their proportion declined after ∼4 ka (Supplementary Fig. 8).

**Fig. 5:**
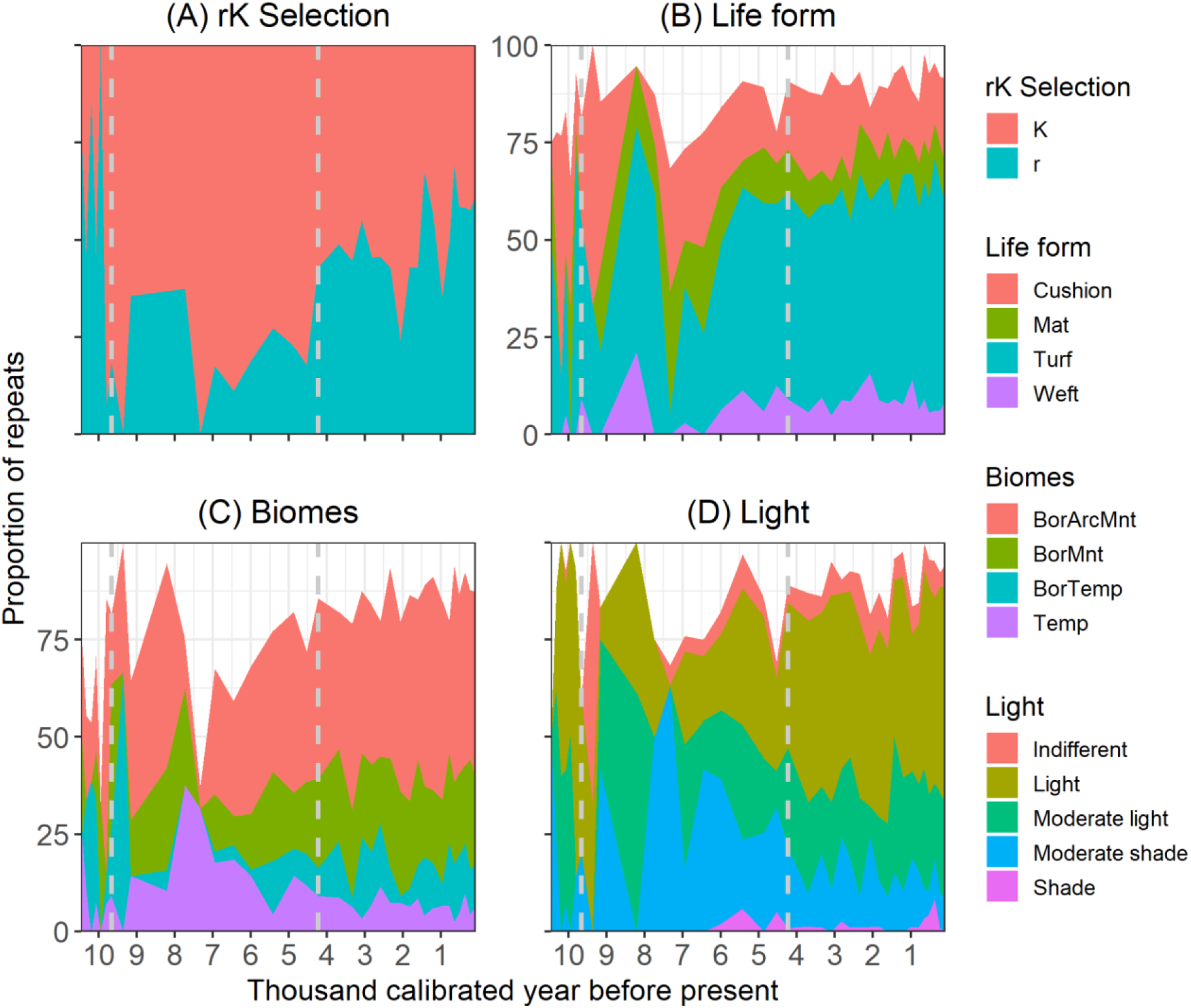
Bryophyte trait variation over time reflected by change in proportion of repeats of different (A) rK selection, (B), Life form, (C) Biomes, and (D) Light. The missing values are indicated by white space. Two vertical dashed lines enclose a period of low glacial activity. Abbreviations for x-axis text: BorArcMnt - Boreo-arctic montane, BorMnt - Boreal-montane, BorTemp - Boreo-temperate, Temp - Temperate.

### Trait-based successional patterns

We observed a high temporal dispersion in the traits, with overlaps in the first arrival date (FAD). The mean value of the FAD indicated a pattern, with pleurocarpous being the earliest arriving taxa, followed by acrocarpous, foliose, thalloid, and *Sphagnum*. Among the growth forms, pleurocarpous, acrocarpous, and foliose were the most abundant traits, with greater variation on a temporal scale (Fig. 6A). Similarly, the mean FAD across life forms indicated that cushion forming species appeared earliest, while all other forms appeared later, but around the same time. Taxa with cushion life forms had higher occurrences during the earlier stages of ecosystem development. Turf was the most frequent trait, with high temporal variation. Turf life form clustered particularly during the early and late Holocene (Supplementary Fig. 9). Though some of the taxa characteristics of temperate biome were present during the early Holocene, most of the early arriving taxa were characteristics of present-day boreo-arctic montane biomes which were the most abundant biomes with larger variation in FAD (Fig. 6B). Taxa with small and medium-sized spores appeared earlier and around the same time; however, those with large spores, though less frequent, appeared much later (Fig. 6C). Taxa preferring open areas appeared during the early Holocene, followed by those occupying both forest and open habitats. However, taxa favoring closed forests emerged later during mid Holocene around 5 ka (Fig. 6D). Bryophyte taxa adapted to light as well as those indifferent to light conditions, appeared earlier, followed by taxa moderately adapted to light, moderately adapted to shade, and finally, shade-adapted taxa (Supplementary Fig. 10).

**Fig. 6:**
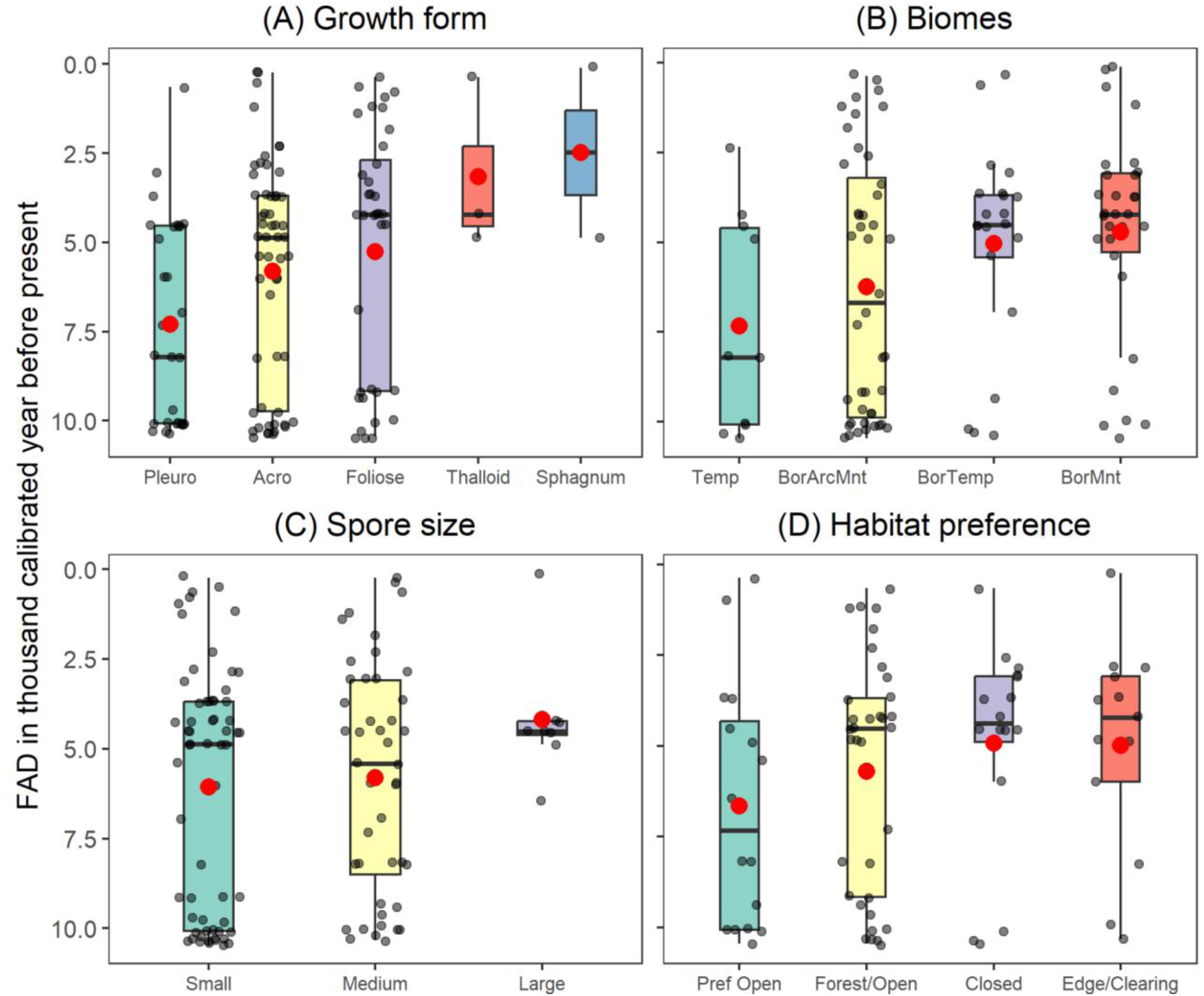
First arrival date of bryophyte taxa with different (A) Growth form, (B) Biomes, (C) Spore size, and (D) Habitat preference in terms of forest canopy. The red dot within the box represents the mean value of the FAD and the zittered dots represent the number of taxa sharing the traits. Abbreviations for x-axis text: Acro - Acrocarpous, Pleuro - Pleurocarpous, WideTemp - Wide temperate, BorArcMn - Boreo-arctic montane, WideBor - Wide-boreal, Temp - Temperate, ArcMnt- Arctic-montane, BorTemp-Boreal-temperate, BorMn - Boreal-montane, Pref Open-may occur in forest, but prefers open land, Forest/Open - occurs in forest as well as in open land, Closed-largely restricted to closed forest, Edge/Clearing - prefers forest edges and clearings

## Discussion

### Bryophyte primers increase taxonomic resolution

Bryophyte-specific primers detected 2.63 times higher bryophyte taxa than bycatch data obtained from vascular plant-specific primers. The effectiveness of the bryophyte-specific primer can be attributed to the targeted amplification of bryophyte DNA. The vascular plant-specific primers, designed to capture a wider range of vascular plant taxa, are less specific to bryophytes as expected, resulting in the predominant amplification of vascular plant sequences and thereby limiting the detection of bryophytes ^9,12^. With preferential amplification, the bryophyte-specific primer primarily allocated reads to bryophyte taxa, leading to improved detection rates. The GC content of the amplicons has been often reported to be associated with PCR success ^39,40^. The GC content in the P6 loop of plants, however, is highly variable ^41^, with lower values in bryophytes (mean±SD = 13.6±7.6%) ^42^. The optimal *trn*L amplification has been observed in vascular plants with a GC content ranging from 34–38% ^39^, which is substantially higher than the GC content found in the P6 loop in bryophytes ^42^. In our case, it is likely that the GC content in the amplicons produced by bryophyte-specific primers may represent an optimal range for these taxa facilitating efficient amplification of bryophytes DNA.

The amplicons generated by the bryophyte-specific primers were considerably longer than bycatch amplicons and provided species-level resolution for nearly 60% of the detected taxa, which is nearly 1.5 times higher than that offered by the bycatch data. Amplicon length plays a key role in taxonomic resolution, with longer amplicons generally providing less ambiguous identifications due to increased alignment coverage against reference genomes ^43–45^. The amplicon length for bryophytes captured in our study is longer than those reported in other *seda*DNA research on bryophytes utilizing the vascular-plant specific primers ^9^, which may have mapped well to the marker region, offering higher taxonomic resolution. The number of bryophyte taxa reported in this study exceeds that of previous *seda*DNA studies ^9,46^. Given the high variation in taxonomic resolution of the *trn*L P6 loop reported across different studies ^9,12,47–49^, the use of bryophyte-specific primers combined with a reference library curated for the local flora can significantly improve identification accuracy and maximize the number of detected taxa ^50^. Therefore, we anticipate further improvements in taxonomic richness and resolution if the reference library includes nearly all bryophyte species.

### Rise of bryophytes diversity in the Holocene and the potential drivers

Temporal richness trends closely resembled glacial activity of the Langfjordjøkelen ice cap, which drains into the Jøkelvatnet. We reported richness increase with glacier activity until a certain threshold, beyond which it declined showing a hump shaped curve. This pattern is consistent with plant successional dynamics observed in glacier forelands, where diversity initially increased with the glacier retreat but declined during the later phase of the community development ^51^. During the period of early high glacial activity (∼10.4 to 9.5 ka) ^27^, we reported low richness but high compositional variation marked by the presence of pioneering and early successional taxa, including *Grimmia* species such as *G. longirostris*, *G. caespiticia*, *G. mollis*, as well as *Scapania subalpina*, *Diphyscium foliosum*. The ecological requirements of earlier bryophyte communities resemble the vascular plant community as they were also represented by a few cold-tolerant taxa belonging to family Salicaceae ^12,13^. During the period of low glacial activities (∼ 9.5-4.0 ka), the bryophyte richness increased almost exponentially. During this time, the catchment was almost completely deglaciated ^27^, probably forming a stable and suitable habitat for further bryophyte community development. Glacier activity affects the accumulation of organic matter, shaping soil profiles, and creating conditions for the establishment of early plant colonizers as well as later successional species, which together drive community diversification ^52,53^. We documented the increased dominance of K-selected taxa during the mid Holocene, low glacier activity period, which may be related to the formation of an environment favorable for the growth of secondary successional taxa. The increased dominance of K selected taxa during mid Holocene aligns with those observed in Siberia and Alaska ^9^. The richness continued to build up even after the onset of glacier activity during the late Holocene, likely utilizing the soil profile and the substrates created earlier; however we also documented the increased dominance of early successional taxa during this time which is consistent with the increased soil disturbance in the catchment following mid Holocene ^13^. The period of exponential increase in bryophyte diversity during the mid Holocene coincides with the buildup of vascular plant richness in the catchment ^13^ suggesting similar responses of both bryophyte and vascular plants to the environmental changes.

Bryophyte richness increased during the Holocene with subtle variations in the magnitude over time. We report nonlinear partial effects of mean precipitation and temperature of the warmest quarter, as well as significant combined effects of mean temperature and precipitation during the warmest quarter. The long-term reports on the effect of temperature and precipitation on bryophytes diversity is fairly lacking, but the short-term, fine-scale reports are inconsistent ^54–57^. In Northern Fennoscandia, a previous study found that temperature positively affected terrestrial plant richness in the early Holocene, but had slight to significant negative effects in the middle and late Holocene, when temperatures were stable and then declined, respectively ^11^. Our finding on a temporal scale indicated that bryophytes may benefit from the combined effect of temperature and precipitation though the effect of temperature may be limiting especially at their higher level. Following deglaciation in the early Holocene, temperatures remained below freezing with low precipitation ^27^. Our observation of low bryophyte richness during this period aligns with the pattern observed during the cold, dry climate and intense glacial activity ^9^. During the mid Holocene, both temperature and precipitation increased ^34^, a combination of conditions that appears to have favored bryophyte diversity, as reflected in the exponential increase in richness. Bryophyte responded well with change in temperature with the change in composition of the taxa adapted to a warmer environment ^9^. Our findings in the scale of millennia suggested that the increase in mean temperature of the warmest quarter limits the bryophyte richness as it declines sharply with temperature after threshold of around 12 ℃, however, increase in precipitation to some extent appears to buffer the negative impact of the temperature. As the Arctic is experiencing warming faster than the global average ^4^, these findings are important to advance our understanding on the potential impact of ongoing climate change to bryophyte diversity in the Arctic.

### Potential for environmental reconstruction

We observed bryophyte traits variability through time, reflecting the response of bryophytes to the changing environment. Sensitivity of bryophytes to environmental change makes them suitable climate change indicators. The high-resolution, species-level taxonomic data of bryophytes can be linked to their traits to study how they responded to past environments at both compositional and functional levels ^58^, thus offering a valuable tool for understanding paleoecology of bryophytes.

There was a higher proportion of r-selected taxa, as expected for a successional community following major disturbance, during the periods of early (before ∼9.5 ka) and late (∼4.0 ka to present) high glacier activities. These taxa were characterized by colonizing species that had an acrocarpous life form and short lifespan. In contrast, the K-selected taxa, which prefer a more stable environment and are competitive in nature ^59^, dominated during the low glacier activities, especially in the mid Holocene. Compositionally, this period formed a distinct bryophyte community with low variability, further suggesting a stable bryophyte community during this phase. The catchment of Jøkelvatnet experienced fluctuating glacial activity with the onset of cooling and warming during the late Holocene, indicating unstable environmental conditions ^27^. Our observation of an increasing abundance of r-selected taxa during this phase likely reflects this environmental instability. Additionally, the proportion of cushion, mat, and turf life forms adapted to dry environments ^60^ also increased during this period, suggesting the onset of dry and cold conditions. The period of increase in abundance of taxa adapted to the dry environments roughly coincides with the built up of glacier activity in the catchment of Jøkelvatnet ^27^.

Though the mid Holocene showed a rise in bryophytes adapted to moderately nutrient-rich environments, we also observed the appearance of *Sphagnum* from the mid to late Holocene, which is adapted to wet and nutrient poor environments ^61,62^. Co-detection of taxa adapted to poor to moderate nutrient conditions and life forms adapted to dry to moist conditions suggests the presence of a complex and diverse environment during the mid Holocene ^23,62^. The mid Holocene also showed an increase in taxa adapted to moderate light and shaded environments, while the late Holocene was again dominated by light-adapted taxa. The detection of taxa growing under the closed forest coincides with an increase in abundance of trees and dwarf shrubs in the catchment during mid Holocene ^13^. Both the early and late Holocene, characterized by high disturbance from glacial activity, exhibited overlapping traits indicative of early community development, whereas the mid Holocene was marked by more stable trait values. The community initially consisted of taxa characteristic of the Arctic environment, represented by *Grimmia mollis*, *Grimmia caespiticia,* and *Scapania subalpina* which increased over time until the present, as also indicated by the rising proportion of taxa from Boreo-Arctic montane biomes. However, the proportion of boreal biomes also increased after the mid Holocene, likely indicating the impact of rising temperatures. Our observation of increasing boreal bryophytes aligns with the findings from Siberia and Alaska ^9^. The dominance of long-lived perennials during the mid Holocene, and colonists during the early and late Holocene, indicates the responses of bryophytes to changing environmental conditions.

As bryophytes occupy narrow niches, this property can be leveraged to infer past ecological changes ^58^. Since *seda*DNA has effectively captured local vegetation shifts and climatic signals in previous studies ^12,13,63^, our finding that bryophyte diversity and composition closely follow local deglaciation, temperature and precipitation patterns, and presence of vascular plants further supports the view that bryophytes are highly sensitive to local-scale environmental changes and can therefore serve as a reliable proxy for reconstructing past environments.

## Conclusion

In the face of a changing global environment and its amplified effects in the Arctic ^3^, there has been a lack of effective molecular tools for generating taxonomically resolved temporal data on bryophytes, which constitute a key component of Arctic biodiversity. Here, we demonstrated the effectiveness of the bryophyte-specific primers targeting the p6 loop of plastid DNA in recovering high-resolution taxonomic information from *seda*DNA. We leveraged this approach not only to investigate past bryophyte diversity but also to assign trait values from a current bryoflora database, allowing us to assess how bryophyte communities functionally responded to past environmental changes.

Our use of bryophyte-specific primers yielded the highest number of taxa from sediment samples, with most resolved to the species level, demonstrating their effectiveness as a metabarcoding tool for studying temporal changes in bryophytes communities using *seda*DNA. These primers also captured the buildup of taxonomic richness over time in ways that aligned well with environmental changes in the catchment. Assigning species-level trait values to majority of taxa enabled functional reconstruction of past bryophyte communities. This highlights the potential of *seda*DNA not only for tracking ecosystem changes but also for examining functional responses of bryophytes to past environmental change. Although our study is limited to a single lake, applying this approach across multiple sites could provide broader insights into bryophyte responses to environmental change at regional and global scales.

## Acknowledgments

ArcEcoGen and the Arctic University of Norway (UiT) for financially supporting the research.

## Authors contribution

**Bishnu Timilsina** laboratory work, bioinformatics, data analysis, visualization, writing-initial draft, writing-review and editing.

**Dilli Prasad Rijal** conceptualization, resource acquisition, planning, methodology, supervision, guided: bioinformatics, data analysis, visualization, writing -review and editing.

