## Supplementary materials for "Enhanced reconstruction of Holocene bryophytes from sedimentary ancient DNA using bryophyte-specific primers"

### Supplementary figures

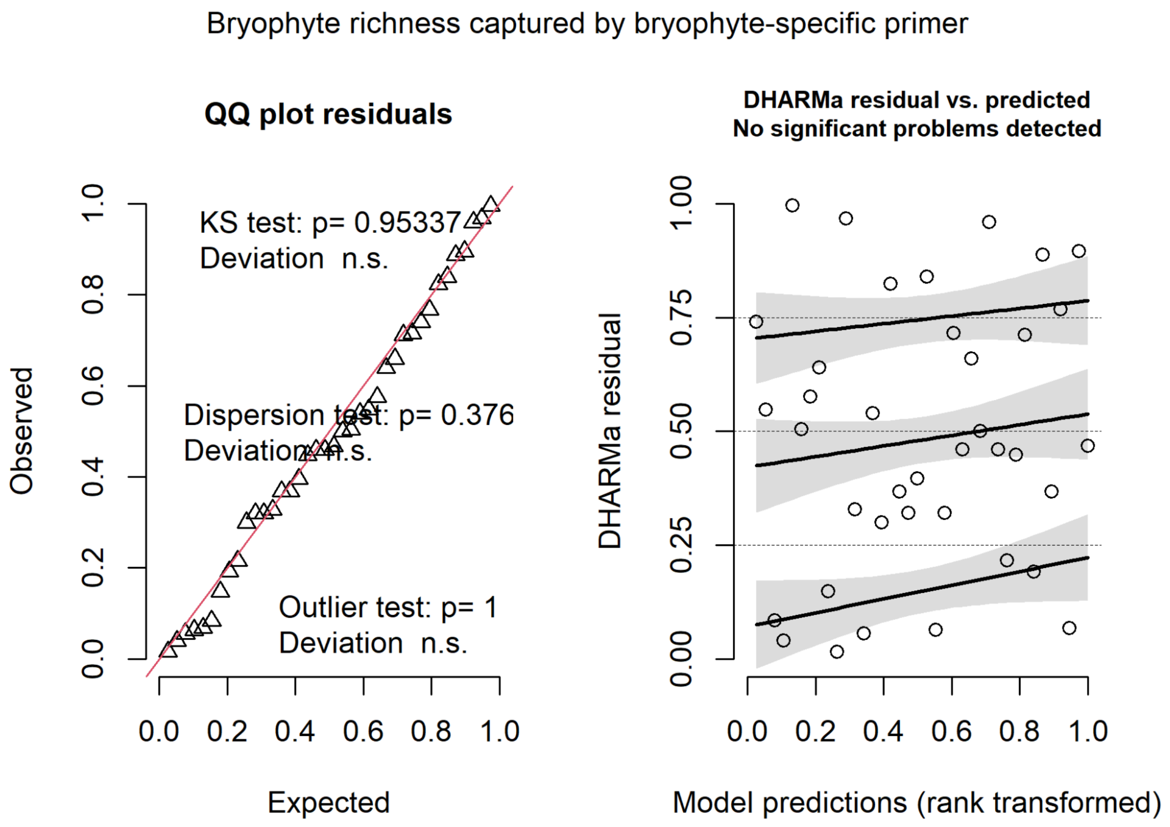


**Supplementary Fig. 1A:** Diagnostic plots indicating fulfillment of model assumptions while evaluating temporal trends of bryophyte richness captured by bryophyte-specific primers.

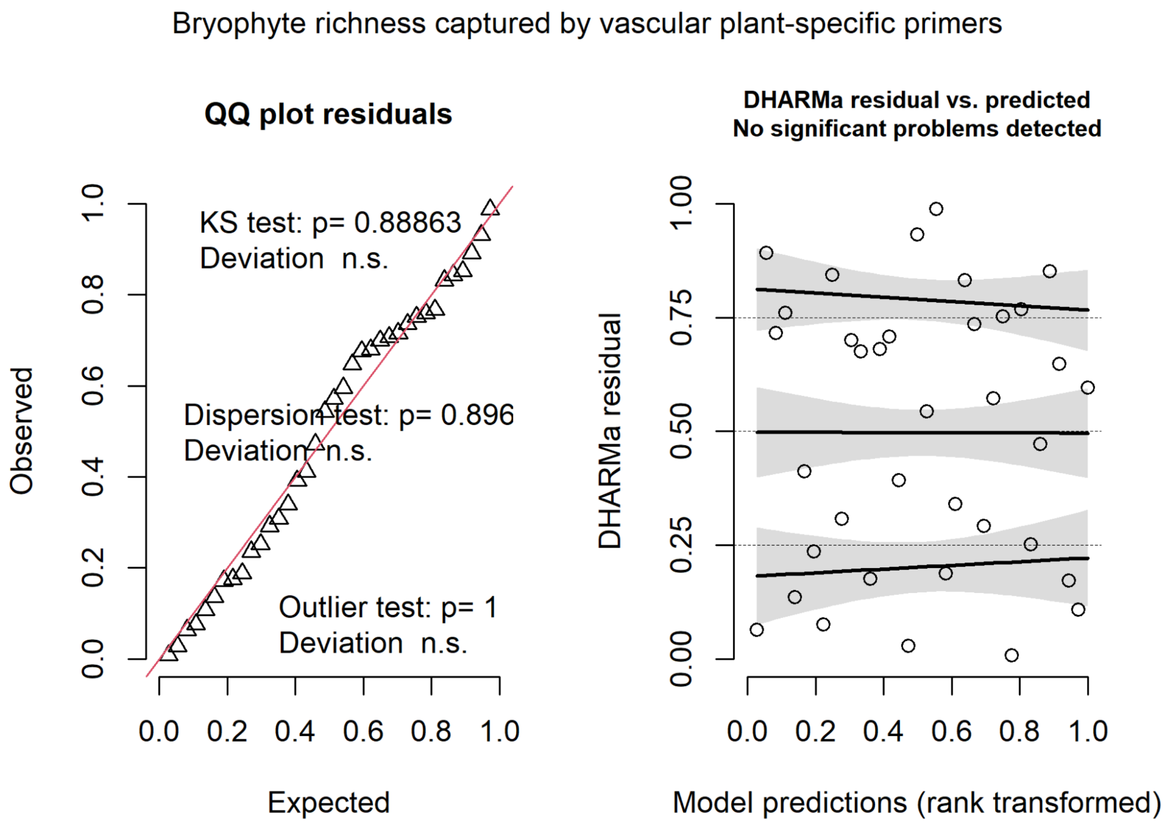


**Supplementary Fig. 1B:** Diagnostic plots indicating fulfillment of model assumptions while evaluating temporal trends of bryophyte richness captured by vascular plant-specific primers.

Impact of glacial activity on bryophyte richness

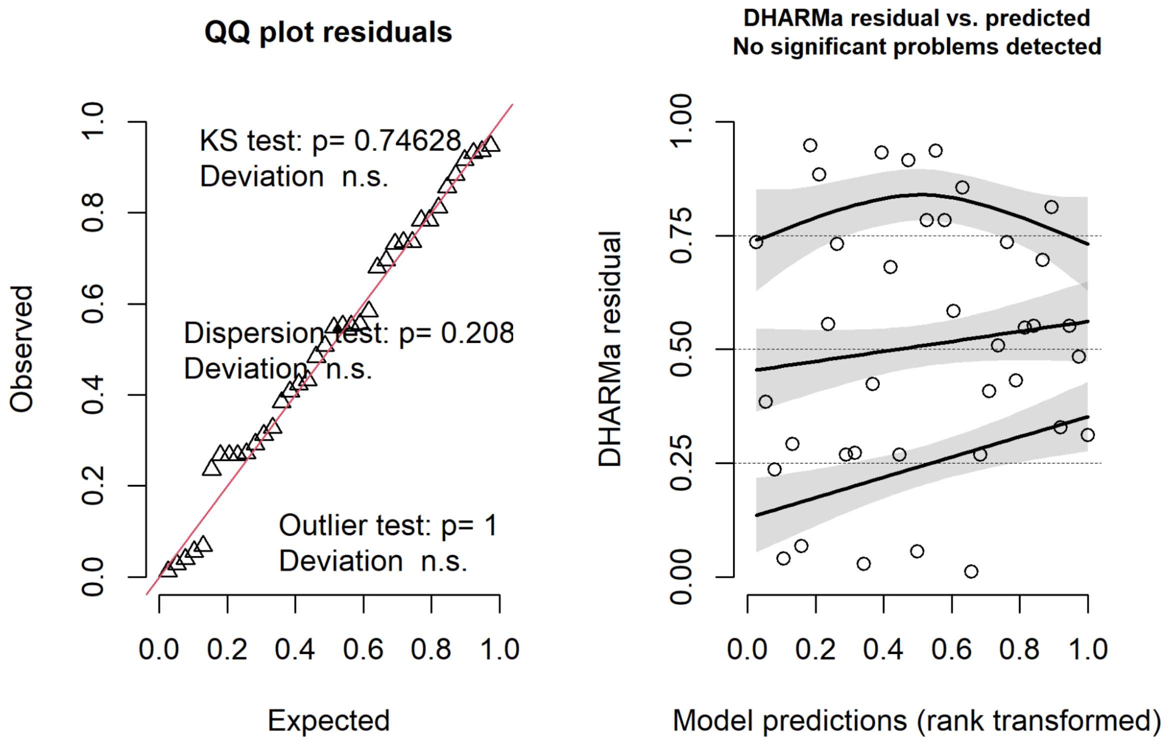


**Supplementary Fig. 2:** Diagnostic plots indicating fulfillment of model assumptions while evaluating impact of glacial activity on bryophyte richness captured by bryophyte-specific primers.

Interactive effect of temperature and precipitation on bryophyte richness

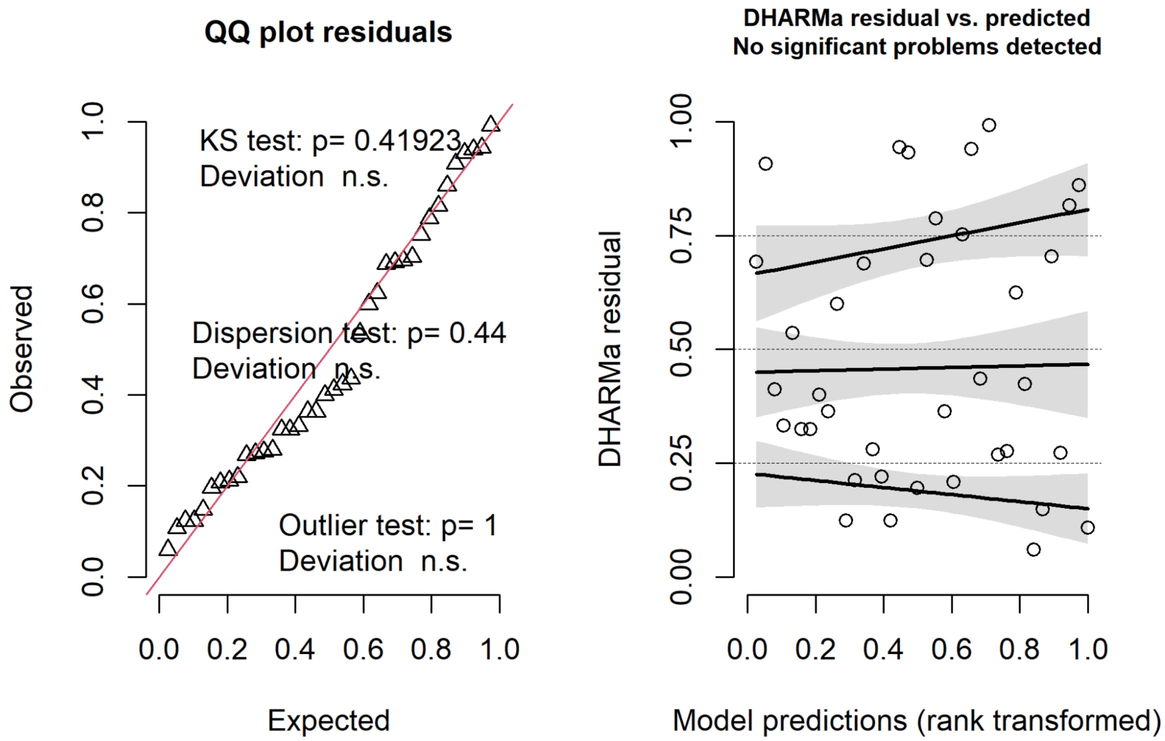


**Supplementary Fig. 3:** Diagnostic plots indicating fulfillment of model assumptions while evaluating the impact of mean temperature and precipitation of the warmest quarter on bryophyte richness captured by bryophyte-specific primers.

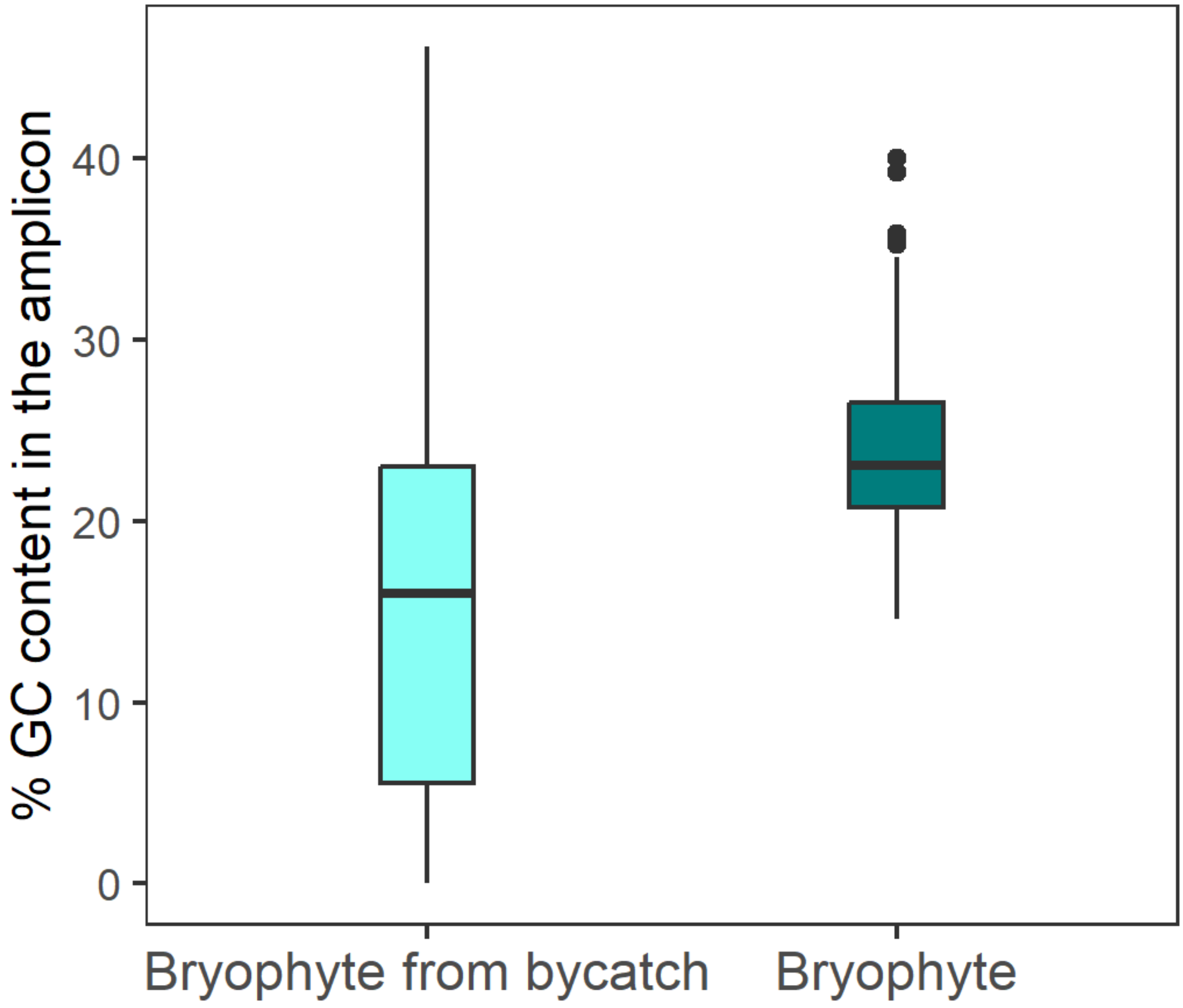


**Supplementary Fig. 4:** Comparison of GC content in the amplicon produced by the vascular plant and the  bryophyte-specific primers. The amplicon produced by the latter has higher GC content.

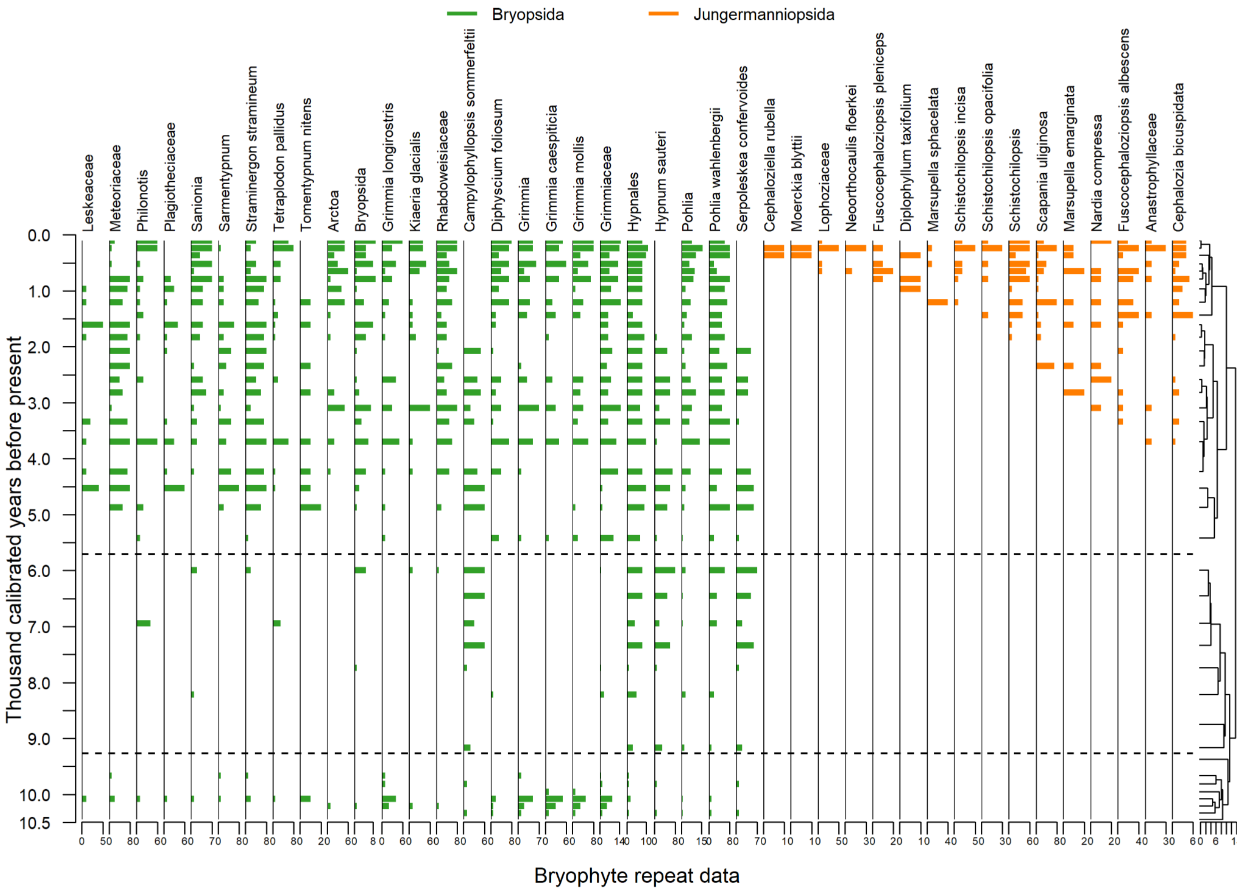


**Supplementary Fig. 5A:** Appearance of bryophyte taxa in Jøkelvatnet. The horizontal dotted lines indicate vegetation zones identified by constrained incremental sum of square (CONISS) analysis on the proportion of PCR repeats. The number of PCR repeats of the taxon in a sample is plotted in the X- axis. Colours differentiate classes of bryophyte.

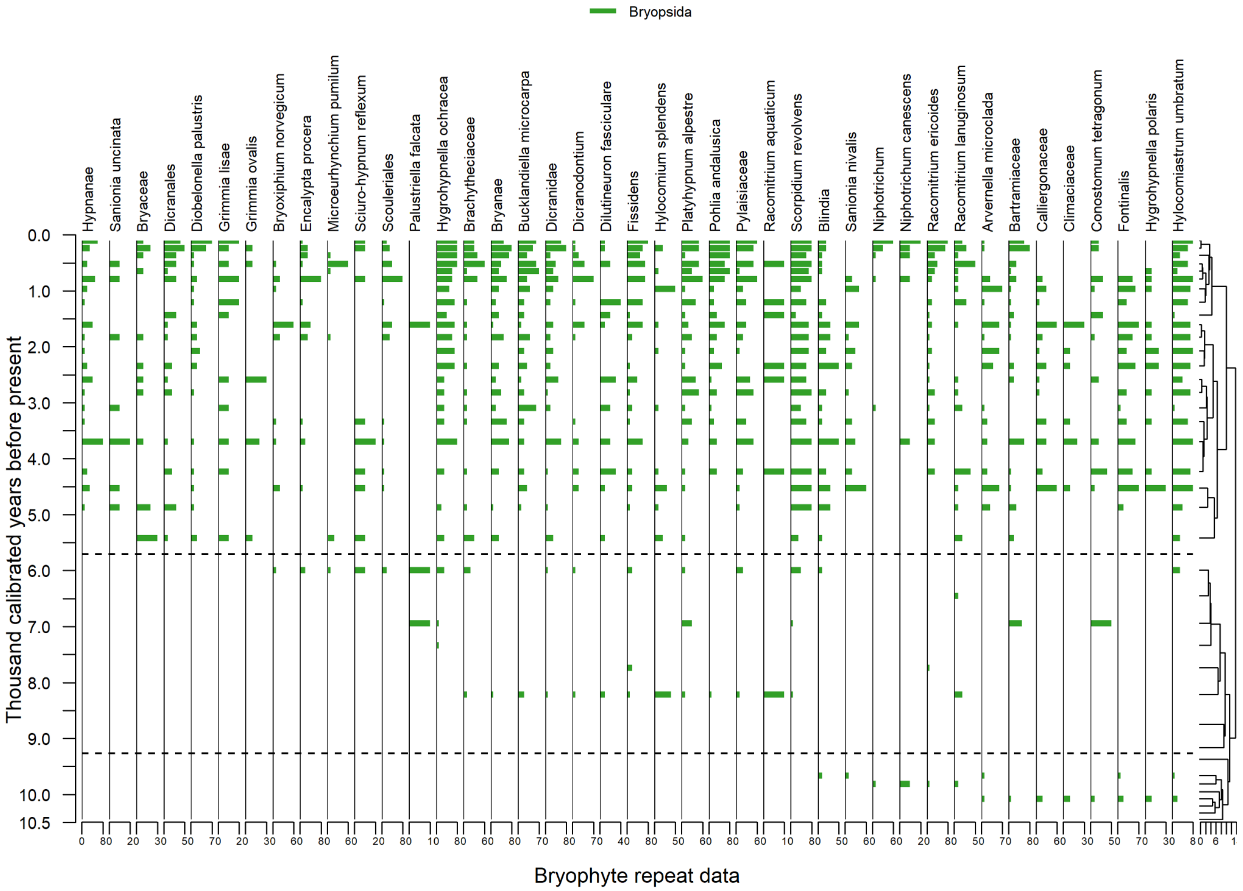


**Supplementary Fig. 5B:** Appearance of bryophyte taxa in Jøkelvatnet. The horizontal dotted lines indicate vegetation zones identified by constrained incremental sum of square (CONISS) analysis on the proportion of PCR repeats. The number of PCR repeats of the taxon in a sample is plotted in the X- axis. Colours differentiate classes of bryophyte.

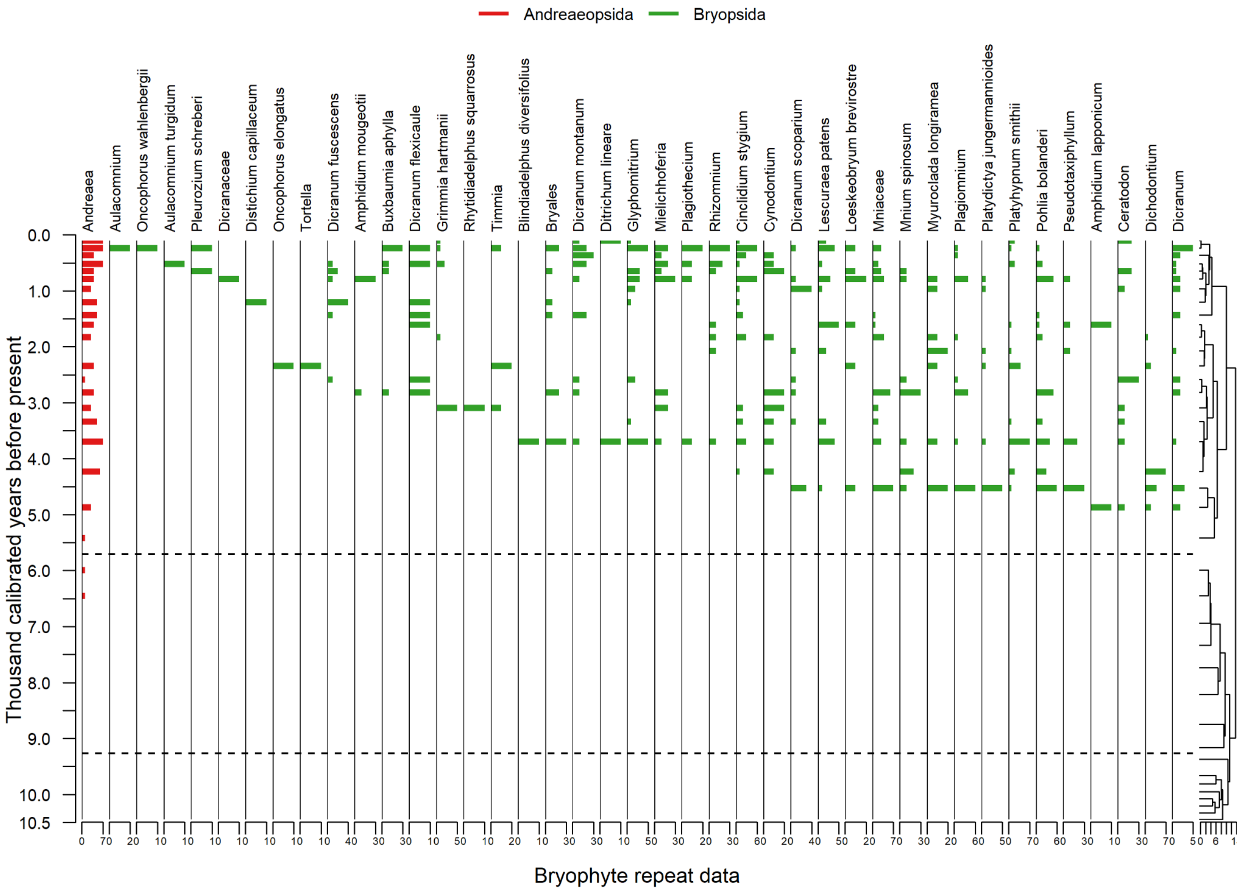


**Supplementary Fig. 5C:** Appearance of bryophyte taxa in Jøkelvatnet. The horizontal dotted lines indicate vegetation zones identified by constrained incremental sum of square (CONISS) analysis on the proportion of PCR repeats. The number of PCR repeats of the taxon in a sample is plotted in the X- axis. Colours differentiate classes of bryophyte.

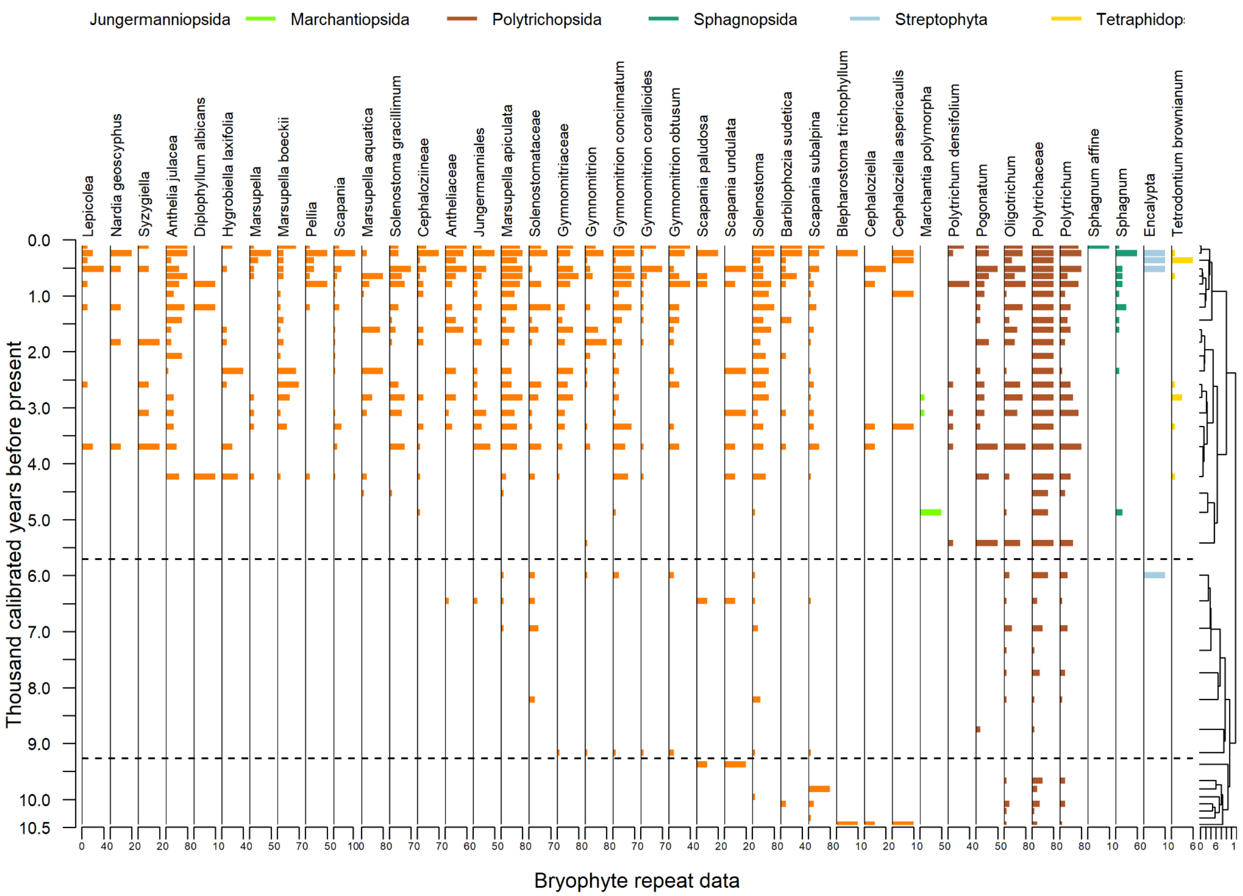


**Supplementary Fig. 5D:** Appearance of bryophyte taxa in Jøkelvatnet. The horizontal dotted lines indicate vegetation zones identified by constrained incremental sum of square (CONISS) analysis on the proportion of PCR repeats. The number of PCR repeats of the taxon in a sample is plotted in the X- axis. Colours differentiate classes of bryophyte.

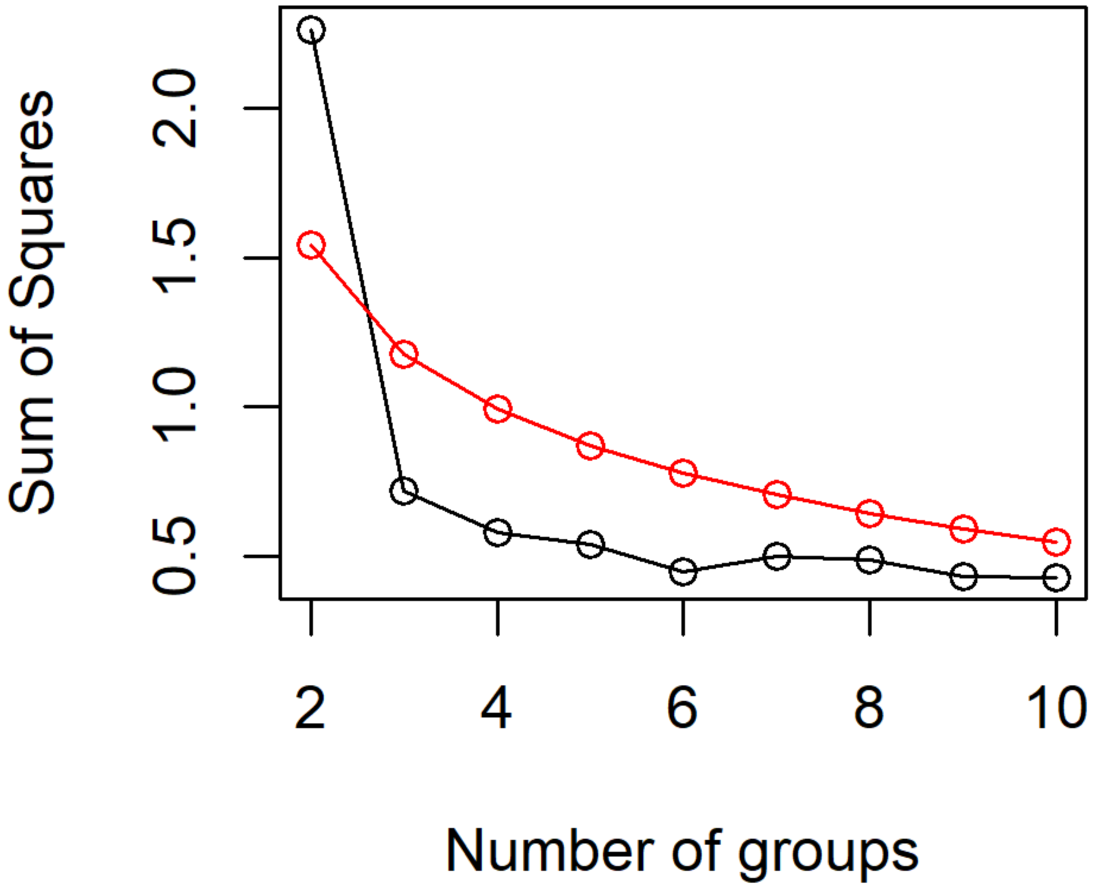


**Supplementary Fig. 6:** Broken stick plot showing actual (black) and theoretical (red) values of the sum of squares for different clusters.

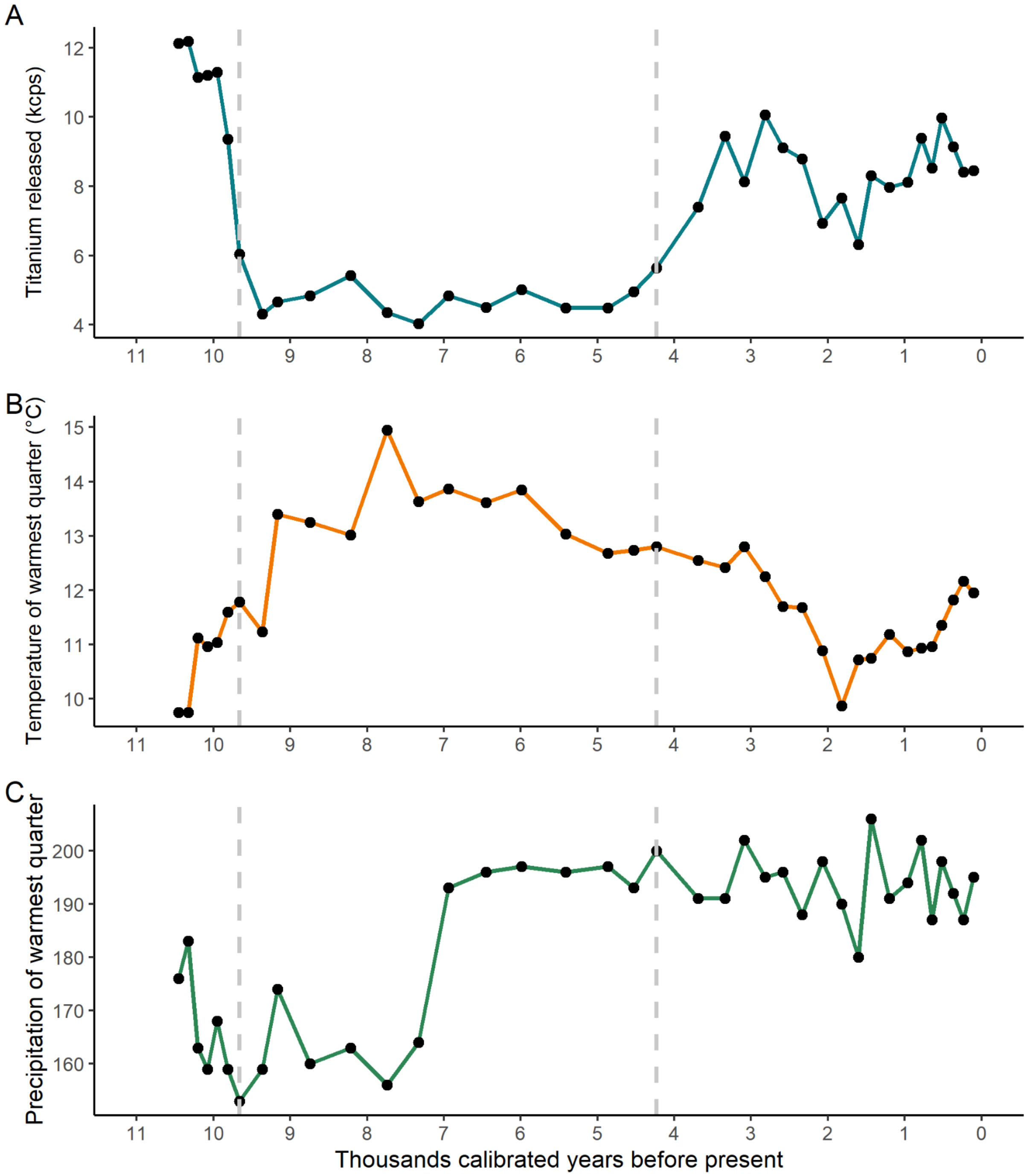


**Supplementary Fig. 7:** Trend of environmental change in Jøkelvatnet lake catchment. A) Glacial Activity measured in terms of titanium release from X-ray fluorescence analysis, B) Mean temperature of the warmest quarter, and C) Mean precipitation of the warmest quarter.

The region between the vertical dotted lines represents the period of low glacial activity.

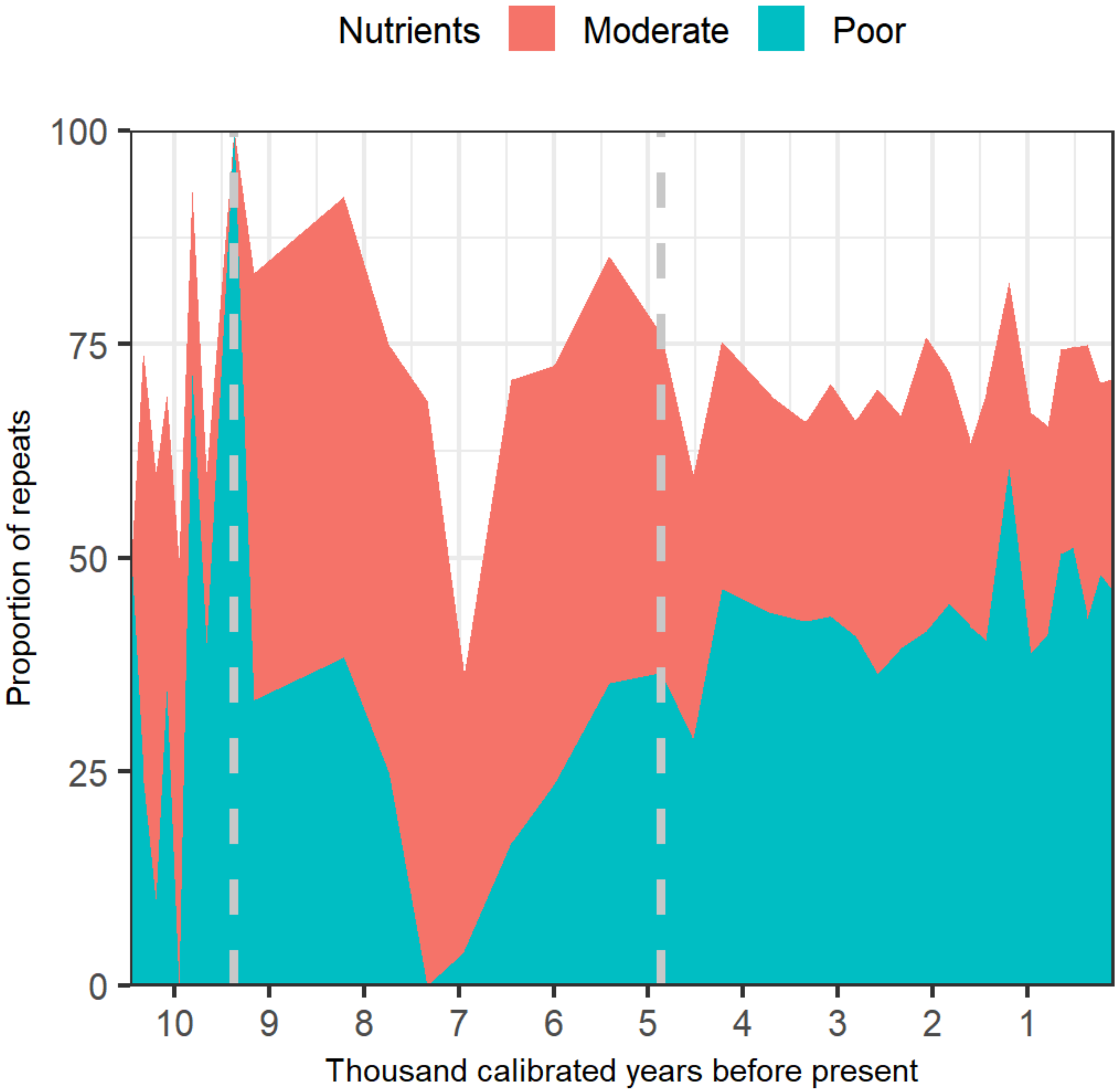


**Supplementary Fig. 8:** Bryophyte nutrient trait variation over time reflected by change in proportion of repeats. The region between the vertical dotted lines represents the period of low glacial activity.

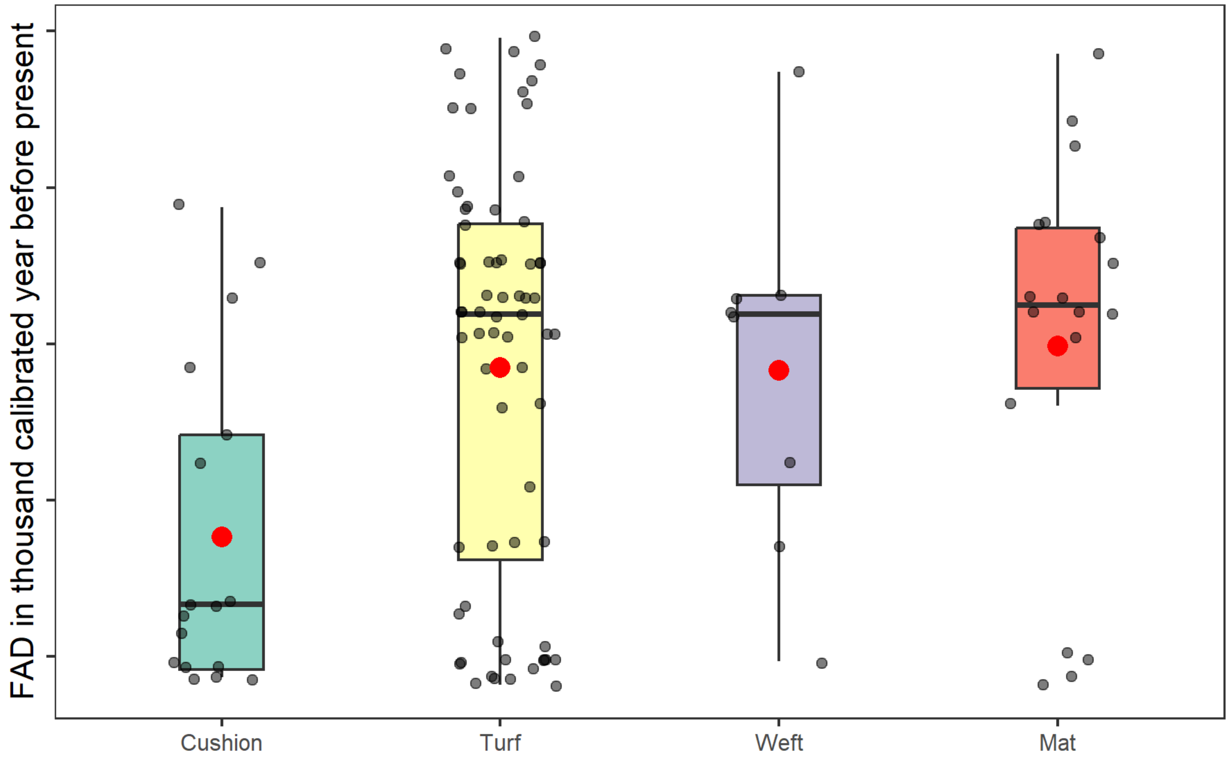


**Supplementary Fig. 9:** First arrival date (FAD) of bryophyte taxa with different life forms. The red dot within the box represents the mean value of the FAD and the zittered dots represent individual FAD.

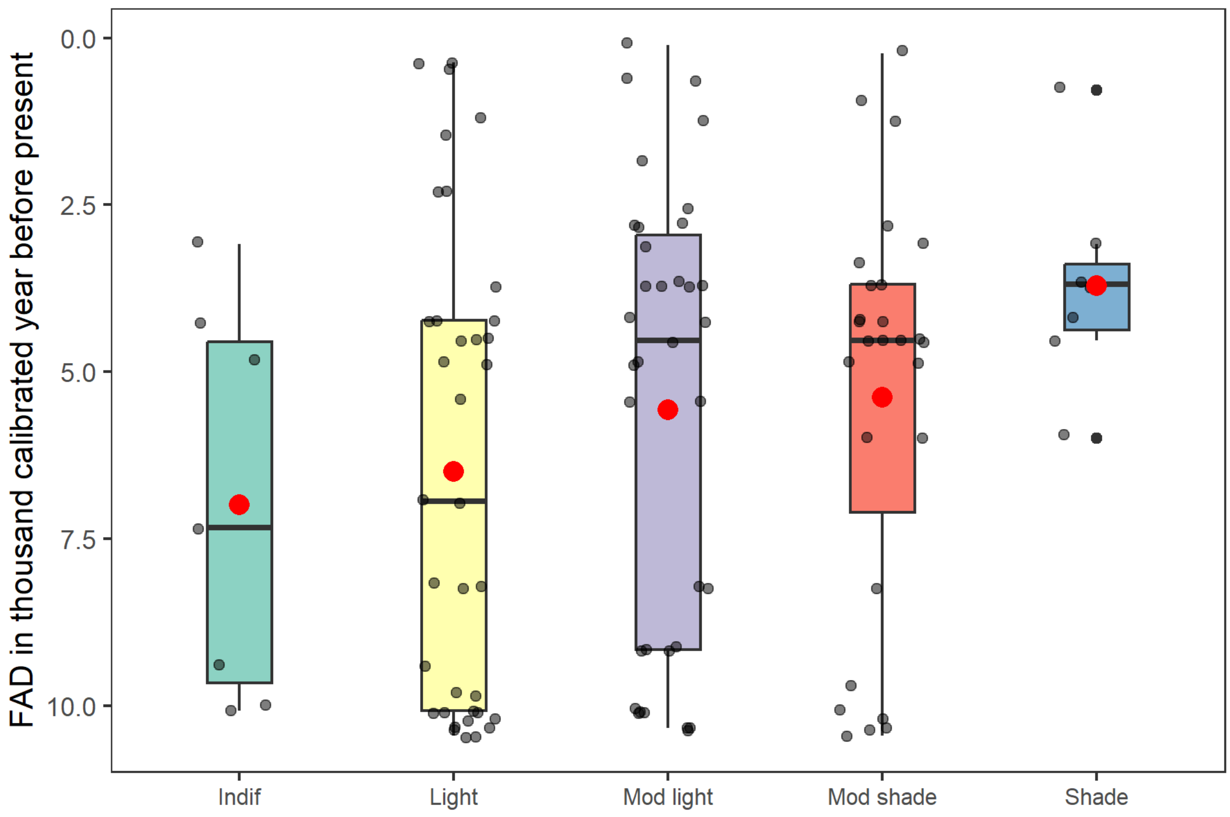


**Supplementary Fig. 10:** First arrival date of bryophyte taxa adapted to different light conditions. The red dot within the box represents the mean value of the FAD and the zittered dots represent individual FAD. Abbreviations for x-axis text: Indif - indifferent, Mod - moderate.

### Supplementary text

**Supp Text:** Bryophyte richness and composition in CONISS identified vegetation zones

The pairwise comparison of taxonomic richness between the two consecutive CONISS zones using Dunn's test revealed differences among the pairs in incremental order over time, though the result was not consistent throughout. There was no significant difference in taxonomic richness between Zone 1 (10.45-9.36 ka) and Zone 2 (9.37 to 5.99 ka), indicating similar richness levels in the earlier periods. However, Zone 3 (5.99 to present) had significantly higher richness compared to both Zone 1 ($p$ = 0.00) and Zone 2 ($p$ = 0.0002), suggesting an increase in richness from earlier periods. We detected 70 taxa during the Early Holocene, which rose by Mid to 112 and to 163 by Late Holocene.

The key bryophytes of each zone and their climatic significances are described below:

Zone-1 (10.45-9.36 ka)

The time between 10453 to 9368 ka falls under this vegetation zone. This time represents the end of glaciation and transition to thermal optimum, with the significant reduction in the glacial activity (Supp Fig: 8). The mean (±SD) number of taxa detected in this zone was 1.75 (±9.01), which is significantly different from Zone 3, but was similar to Zone 2. *Scapania subalpina* was the most abundant taxa in this cluster. Besides, species such as *Grimmia caespiticia* and *Grimmia mollis* (Supplementary Fig. 4A) also known as the cushion of the bryophyte were also dominant.

Zone-2

The time between 9367 to 5991 ka falls under this zone. This period aligns with the Holocene Thermal Optimum (Supplementary Fig. 6B) and almost no glacier activity (Supplementary Fig. 6A). We detected 17 (± 11.57) taxa per sample in this vegetation zone, which is comparable to Zone 1 but significantly different from Zone 3.

Zone-3

Between approximately 5990 to present ka, bryophyte taxa characterized a distinct vegetation zone. This period overlaps with the latter phase of the Holocene thermal maximum marked by elevated global temperatures. Notably, during the later part of this interval, increased glacial activity and a subsequent decline in temperatures were observed. These climatic shifts influenced the composition and distribution of bryophyte communities. We detected 77.54 (±

20.53) taxa per sample which is significantly different from Zone 1 and Zone 2. Most of the abundant taxa detected during this time, for example *Hylocomiastrum umbratum, Hypnum sauteri, Pohlia wahlenbergii, Scorpidium revolvens, Straminergon stramineum* were adapted to a more or less stable environment.
